# Inclusion of *Salicornia ramosissima* biomass in diets for juvenile whiteleg shrimp (*Penaeus vannamei*) induces favourable but transient effects in the immune and oxidative status

**DOI:** 10.1101/2024.07.05.602095

**Authors:** Lourenço Ramos-Pinto, Raquel Marçal, André Barreto, Adriana Laranjeira, Marina Machado, Sérgio Fernández-Boo, Carla Teixeira, Joana Oliveira, Ana Couto, Jorge Dias, Sofia Guilherme, Ana C. S. Veríssimo, Diana C. G. A. Pinto, Mário Pacheco, Rui J.M. Rocha, Benjamín Costas

## Abstract

The whiteleg shrimp, *Penaeus vannamei*, is a highly valued and globally produced crustacean species. However, the rising cost of shrimp feed, exacerbated by increasing cereal prices, prompts the exploration of cost-effective and sustainable formulations for shrimp farming. This study investigates the potential of *Salicornia ramosissima*, a non-edible biomass co-product, as a substitute for wheat meal in juvenile whiteleg shrimp diets, aiming to create economically and ecologically sound formulations. The present study aimed to assess the impact of incorporating *S. ramosissima* into whiteleg shrimp aquafeeds on various aspects of shrimp development, including growth performance, survival, immune status, and oxidative status. A commercial-like diet was formulated and served as control, whereas four other diets contained *S. ramosissima* stems or a combination of leaves and seeds, both at inclusion levels of 5% and 10%, in addition to the control diet. Whiteleg shrimps were fed the experimental diets for 31 and 55 days, followed by a bacterial bath challenge to gauge their immune response to pathogens.

At the end of the feeding period, whiteleg shrimps’ growth performance and survival rates remained consistent across all diets. However, whiteleg shrimp fed diets with *S. ramosissima* inclusion consumed more feed to achieve similar weights to those fed on the control diet, particularly in diets containing leaves and seeds at a 10% inclusion level, likely due to lower digestibility of dry matter, lipids, and energy. While *S. ramosissima* biomass inclusion did not affect shrimp weight, relative growth rate, or survival, it did lead to higher feed conversion ratios and feed intake, suggesting differences in nutrient digestibility and metabolic utilization. Additionally, *S. ramosissima* inclusion affected whiteleg shrimps’ overall body composition, particularly moisture and ash content. Dietary *S. ramosissima* inclusion modulated antioxidant enzyme activity in the shrimp’s hepatopancreas, indicating potential health improvements. The study also observed gene expression changes related to antioxidant enzymes, indicating an overall down-regulation with the inclusion of *S. ramosissima*. Despite challenges in feeding efficiency, the inclusion of *S. ramosissima*, especially stems, shows promise in reducing feed costs by utilizing a by-product. Furthermore, *S. ramosissima* inclusion led to subtle changes in certain plasma humoral parameters and hepatopancreas gene expression. Although some immune parameters varied, these effects appeared to diminish over time. In conclusion, this study highlights the potential of *S. ramosissima* as a functional feed ingredient capable of enhancing shrimp’s antioxidant response, aligning with global resource optimization and sustainability initiatives.

## 1. Introduction

The increasing challenge of soil salinization is a significant constraint on agricultural productivity globally. About 7 % of the world’s land area is salt-affected to some degree, and this trend is on the rise ^1^. Glycophytes, which are the primary source of food, home remedies, and drugs, are sensitive to salt and suffer yield reductions even in mildly saline conditions. Halophytes, on the other hand, have been identified as a viable alternative in recent decades. These plants are adapted to grow in saline environments, such as coastal areas and salt marshes, and can tolerate high salt concentrations of 200 mM or more in their tissues, surviving in conditions that most other plants cannot. They make up about 2 % of the world’s flora ^2,3^. However, it is only recently that their potential has been recognized for crop production, as they do not require as much freshwater for irrigation and can tolerate high levels of salt and other environmental stresses ^4^. Halophytes have a variety of applications, including food, medicine, and biofuels. Species of the genus *Salicornia* are particularly noteworthy due to their commercial potential, with current large-scale cultivation in Europe, North America and middle East for human consumption as a vegetable, as well as attempts to produce oilseed and animal feed using seawater irrigation ^5^.

The increasing interest and widespread embrace of functional foods have paved the way for a fresh market opportunity for halophytes. This stems from the revelation of various bioactive compounds found in halophytes, which not only contribute to their nutritional value but also assist in specific bodily functions. Research indicates that *Salicornia* spp. possess numerous nutraceutical properties attributed to their phenolic content ^6^. They have been used traditionally in the treatment of conditions such as hepatitis, constipation, nephropathy, diarrhea, and have demonstrated antihyperglycemic and antihyperlipidemic activities. Some of the phenolic compounds in *Salicornia* spp. are recognized as generally safe by the United States Food and Drug Administration (USFDA) and can be administered in specific concentrations. The concentrations of phenolic compounds in *Salicornia* spp. can vary due to factors like biomass maturity, pre-treatment methods, and extraction techniques. Sub-critical water extraction and deep eutectic solvents have emerged as promising methods for extracting phenolic compounds from *Salicornia* spp.. Additionally, resin adsorption and desorption techniques have shown promise in concentrating phenolic compounds. In terms of antiviral properties, phenolic compounds found in *Salicornia* spp. have demonstrated inhibitory effects against viruses such as H1N1, HBV, HCV, and HIV-1. These effects are attributed to the compounds’ ability to interact with viral and host proteins and enzymes, preventing viral replication and infectivity^6^. Hence, functional feeds – such as diets rich in *Salicornia* – may play a crucial role in enhancing the overall health and performance of aquatic organisms in aquaculture, particularly in species like shrimp. By incorporating functional ingredients into their diets, it is possible to boost their immune system and improve antioxidant functions. This could help the shrimp to better combat diseases and oxidative stress, ultimately leading to improved growth, survival rates, and overall welfare. This is particularly vital in aquaculture, where maintaining the health and vitality of the stock is essential for successful and sustainable production.

*Salicornia ramosissima* is considered an alternative to salt and its fresh and tender green tips (leaves) are used as food ingredients for human diet, mainly in salads, canned or gourmet gastronomy ^7^. However, as the plant grows, it becomes woody and unappetizing due to lignification and high salt content. Recent research has shown that *Salicornia* spp. can grow in constructed wetlands and can effectively take up nutrients, such as nitrogen and phosphorus, from wastewater into its tissues ^4^. *Salicornia spp.* are rich in minerals ^8^, flavonoids and polyphenols ^9-11^. Polyphenols have been found to be strong antioxidants that can neutralize free radicals by donating an electron or hydrogen atom ^12^. Flavonoids possess antioxidant activity and function as a free radical scavenger and a chelator of transition metal, such as iron, a key metal in the generation of ROS through the Fenton reaction ^13^. The chemical research on various *Salicornia spp.*, such as *S. europaea*, *S. arthrocnemum*, *S. bigelovii*, *S. perennis*, and *S. disarticulata*, has demonstrated their significant anti-oxidative properties. This finding supports their traditional use, as they all contain phenolic compounds ^14^. Notably, Silva, et al. ^11^ highlighted that *S. ramosissima* surpasses other species in terms of phenolic and flavonoid contents. These compounds derive their antioxidant capabilities from multiple mechanisms, including the scavenging of free radicals ^15^, and the quenching of singlet-oxygen atom ^16^.

The whiteleg shrimp is currently the most produced animal species in aquaculture and over 33 billion USD per year are generated worldwide from its culture ^17^. To sustain the fast growth the industry has experienced, the quantities of aquafeeds used increased accordingly. Still, these usually constitute upwards of 50 % of the overall inputs of production and rely on conventional ingredients, such as fish meal, soybean meal and wheat meal, that raise concerns regarding their sustainability ^17^. Hence, the success of shrimp farming, and aquaculture in general, depends on the emergence of novel ingredients that can make formulated diets more economical and reduce the industry’s ecological footprint.

In summary, this study aimed to explore the viability of integrating *S. ramosissima* biomass into inert diets for juvenile whiteleg shrimp. Additionally, it sought to examine the potential advantages of this inclusion as a functional feed, potentially leading to positive impacts on health and welfare. The assessment encompassed parameters such as growth performance, survival rate, overall health, and oxidative status of the shrimp.

## 2. Material and Methods

### 2.1. Diets formulation

Five experimental diets were evaluated in quintuplicates. A control diet (CTRL) was formulated to meet the nutritional requirements of juvenile whiteleg shrimp containing 32.5 % of soybean meal, 25 % of wheat meal and 15% of fish meal as main protein sources, whereas a blend of 6 % of rice bran oil, 2 % soybean oil and 1 % fish oil was regarded as main lipid source. On the remaining treatments, four experimental variants based on the CTRL were used, differing only in the ingredient formulation by replacing wheat meal by: 1) *S. ramosissima* stems in 5 % (SS5) and 10 % (SS10) inclusion levels; and 2) *S. ramosissima* leaves and seeds in 5 % (SL5) and 10 % (SL10) inclusion levels. *S. ramosissima* biomass was obtained in Praia da Areia Branca, Torreira, Portugal (40°46’22.1”N 8°39’29.5”W). Plants were harvested in September 2019, when signs of lignification and flowering were evident, to simulate the stage of production when *S. ramosissima* is considered inadequate for human consumption. These were then dried in mesh mats that allowed air circulation, to prevent any degradation promoted by humidity. Plant separation occurred naturally in the drying process. The experimental diets composition and proximate analysis can be seen in Tables 1 and 2, respectively.

**Table 1.**
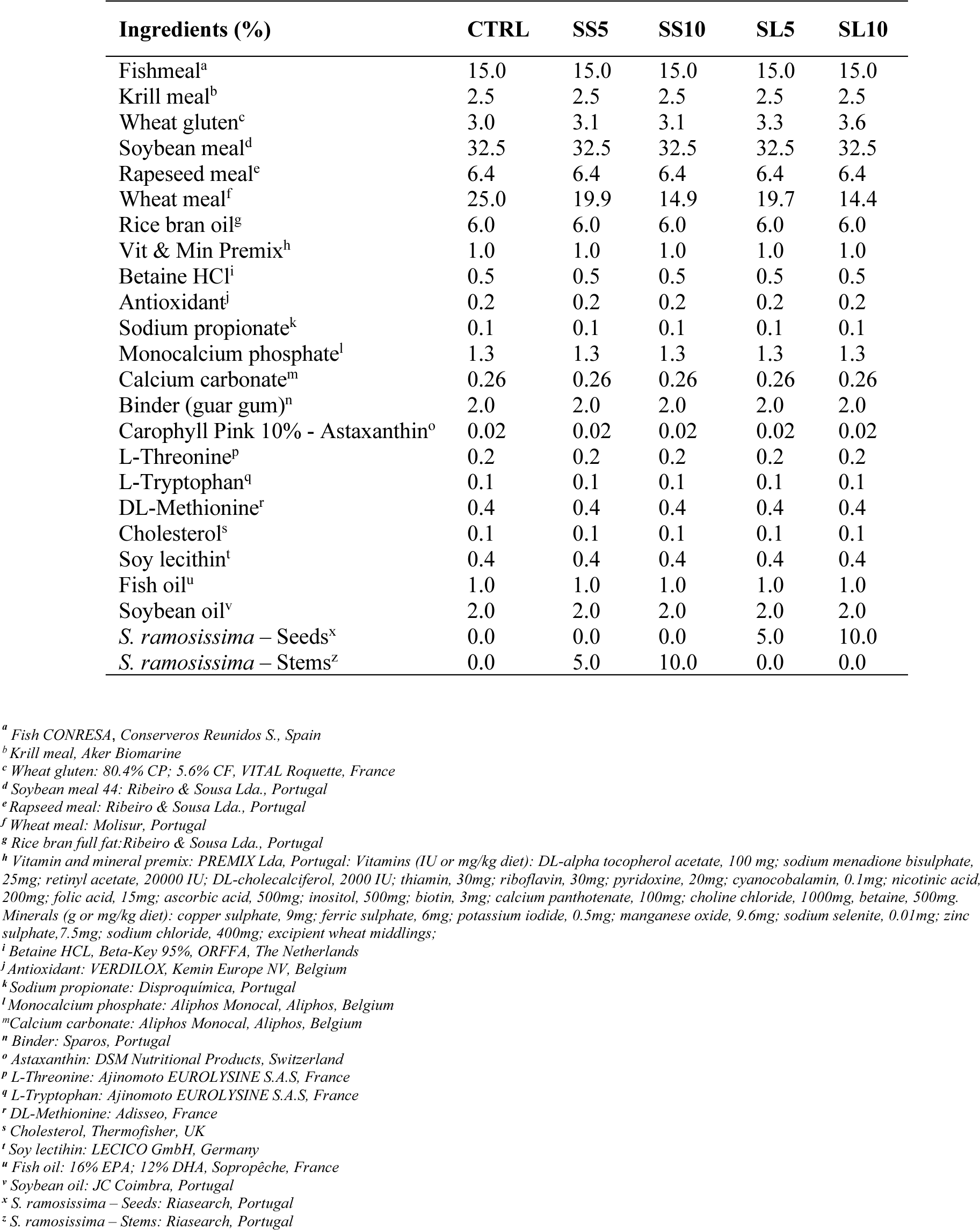
Dietary composition of the experimental diets used to culture juvenile whiteleg shrimp (*P. vannamei*) for 55 days.

| Ingredients (%) | CTRL | SS5 | SS10 | SL5 | SL10 |
| --- | --- | --- | --- | --- | --- |
| Fishmeal<sup>a</sup> | 15.0 | 15.0 | 15.0 | 15.0 | 15.0 |
| Krill meal<sup>b</sup> | 2.5 | 2.5 | 2.5 | 2.5 | 2.5 |
| Wheat gluten<sup>c</sup> | 3.0 | 3.1 | 3.1 | 3.3 | 3.6 |
| Soybean meal<sup>d</sup> | 32.5 | 32.5 | 32.5 | 32.5 | 32.5 |
| Rapeseed meal<sup>e</sup> | 6.4 | 6.4 | 6.4 | 6.4 | 6.4 |
| Wheat meal<sup>f</sup> | 25.0 | 19.9 | 14.9 | 19.7 | 14.4 |
| Rice bran oil<sup>g</sup> | 6.0 | 6.0 | 6.0 | 6.0 | 6.0 |
| Vit & Min Premix<sup>h</sup> | 1.0 | 1.0 | 1.0 | 1.0 | 1.0 |
| Betaine HCl<sup>i</sup> | 0.5 | 0.5 | 0.5 | 0.5 | 0.5 |
| Antioxidant<sup>j</sup> | 0.2 | 0.2 | 0.2 | 0.2 | 0.2 |
| Sodium propionate<sup>k</sup> | 0.1 | 0.1 | 0.1 | 0.1 | 0.1 |
| Monocalcium phosphate<sup>l</sup> | 1.3 | 1.3 | 1.3 | 1.3 | 1.3 |
| Calcium carbonate<sup>m</sup> | 0.26 | 0.26 | 0.26 | 0.26 | 0.26 |
| Binder (guar gum)<sup>n</sup> | 2.0 | 2.0 | 2.0 | 2.0 | 2.0 |
| Carophyll Pink 10% - Astaxanthin<sup>o</sup> | 0.02 | 0.02 | 0.02 | 0.02 | 0.02 |
| L-Threonine<sup>p</sup> | 0.2 | 0.2 | 0.2 | 0.2 | 0.2 |
| L-Tryptophan<sup>q</sup> | 0.1 | 0.1 | 0.1 | 0.1 | 0.1 |
| DL-Methionine<sup>r</sup> | 0.4 | 0.4 | 0.4 | 0.4 | 0.4 |
| Cholesterol<sup>s</sup> | 0.1 | 0.1 | 0.1 | 0.1 | 0.1 |
| Soy lecithin<sup>t</sup> | 0.4 | 0.4 | 0.4 | 0.4 | 0.4 |
| Fish oil<sup>u</sup> | 1.0 | 1.0 | 1.0 | 1.0 | 1.0 |
| Soybean oil<sup>v</sup> | 2.0 | 2.0 | 2.0 | 2.0 | 2.0 |
| *S. ramosissima* – Seeds<sup>x</sup> | 0.0 | 0.0 | 0.0 | 5.0 | 10.0 |
| *S. ramosissima* – Stems<sup>z</sup> | 0.0 | 5.0 | 10.0 | 0.0 | 0.0 |

<sup>a</sup> Fish CONRESA, Conserveros Reunidos S., Spain
<sup>b</sup> Krill meal, Aker Biomarine
<sup>c</sup> Wheat gluten: 80.4% CP; 5.6% CF, VITAL Roquette, France
<sup>d</sup> Soybean meal 44: Ribeiro & Sousa Lda., Portugal
<sup>e</sup> Rapeseed meal: Ribeiro & Sousa Lda., Portugal
<sup>f</sup> Wheat meal: Molisur, Portugal
<sup>g</sup> Rice bran full fat: Ribeiro & Sousa Lda., Portugal
<sup>h</sup> Vitamin and mineral premix: PREMIX Lda, Portugal: Vitamins (IU or mg/kg diet): DL-alpha tocopherol acetate, 100 mg; sodium menadione bisulphate, 25mg; retinyl acetate, 20000 IU; DL-cholecalciferol, 2000 IU; thiamin, 30mg; riboflavin, 30mg; pyridoxine, 20mg; cyanocobalamin, 0.1mg; nicotinic acid, 200mg; folic acid, 15mg; ascorbic acid, 500mg; inositol, 500mg; biotin, 3mg; calcium panthotenate, 100mg; choline chloride, 1000mg; betaine, 500mg. Minerals (g or mg/kg diet): copper sulphate, 9mg; ferric sulphate, 6mg; potassium iodide, 0.5mg; manganese oxide, 9.6mg; sodium selenite, 0.01mg; zinc sulphate, 7.5mg; sodium chloride, 400mg; excipient wheat middlings;
<sup>i</sup> Betaine HCL, Beta-Key 95%, ORFFA, The Netherlands
<sup>j</sup> Antioxidant: VERDILOX, Kemin Europe NV, Belgium
<sup>k</sup> Sodium propionate: Disproquímica, Portugal
<sup>l</sup> Monocalcium phosphate: Aliphos Monocal, Aliphos, Belgium
<sup>m</sup> Calcium carbonate: Aliphos Monocal, Aliphos, Belgium
<sup>n</sup> Binder: Sparos, Portugal

**Table 2.** Proximate composition of the experimental diets used to culture juvenile whiteleg shrimp (*P. vannamei*) for 55 days.

| Proximate composition (% dry matter) | CTRL | SS5 | SS10 | SL5 | SL10 |
| --- | --- | --- | --- | --- | --- |
| Dry matter | 94.8 ± 0.2 | 94.7 ± 0.4 | 93.0 ± 0.0 | 93.7 ± 0.1 | 92.9 ± 0.1 |
| Crude protein | 38.6 ± 0.0 | 38.5 ± 0.3 | 38.3 ± 0.1 | 39.0 ± 0.5 | 39.3 ± 0.2 |
| Crude lipids | 7.9 ± 0.1 | 7.9 ± 0.1 | 7.6 ± 0.1 | 7.9 ± 0.1 | 8.2 ± 0.0 |
| Ash | 8.2 ± 0.6 | 9.9 ± 0.0 | 11.6 ± 0.1 | 10.9 ± 0.7 | 12.6 ± 0.3 |
| Energy (KJ g<sup>-1</sup> DM) | 21.1 ± 0.1 | 20.8 ± 0.9 | 20.3 ± 0.0 | 20.7 ± 0.6 | 20.7 ± 0.2 |

Results expressed as mean ± standard deviation. (n = 3 experimental units).

Diets were manufactured by extrusion at Sparos Lda (Olhão) facilities. All powder ingredients, including *Salicornia*, were mixed accordingly to the target formulation in a double-helix mixer (model 500 L, TGC Extrusion) and ground (below 400 µm) in a micropulveriser hammer mill (model SH1, Hosokawa-Alpine). Diets (pellet size: 1.2 and 2.0 mm) were manufactured with a twin-screw extruder (model BC45, Clextral) with a screw diameter of 55 mm. Extrusion conditions: feeder rate (80 kg/h), screw speed (255 rpm), water addition in barrel 1 (340 ml/min), temperature barrel 1 (36 °C) and temperature barrel 3 (110–112 °C). Extruded pellets were dried in a vibrating fluid bed dryer (model DR100, TGC Extrusion). After cooling, oils were added by vacuum coating (model PG-10VCLAB, Dinnissen). Coating conditions were pressure (700 mbar), spraying time under vacuum (approximately 90 s) and return to atmospheric pressure (120 s). After coating, diets were packed in sealed plastic buckets and stored at room temperature.

### 2.2. Shrimp rearing conditions and samplings

Whiteleg shrimp post-larvae (PL16), originated from Sea Products Development (Texas, USA), were reared for 55 days in standard conditions at Riasearch Lda. facilities (Murtosa, Portugal). After reaching approximately 6 g of body weight, shrimp were weighed individually and randomly distributed to 25 tanks with 200 L, part of a 30 m^3^ clear recirculating aquaculture system (RAS). Each tank was stocked with 55 individuals. Water renewal in each tank was kept at 1 renewal per hour. Water parameters were measured daily using commercial probes. During the experimental period, temperature was maintained at 28.1 ± 0.5 °C, dissolved oxygen at 5.9 ± 0.3 mg L^-1^, salinity at 19.4 ± 0.1 g L^-1^, pH at 7.6 ± 0.1, NH_3_ at 0.0 ± 0.0 mg L^-1^ and NO_2_ at 0.1 ± 0.0 mg L^-1^.

Shrimp were kept under a 12 hours light:12 hours dark photoperiod and given 4 meals per day. Initial feed amounts were established according to Riasearch feeding tables for whiteleg shrimp juveniles. Afterwards, amounts were adjusted daily according to the feed wasted the day before, in increments or decrements of 0, 10, 20 or 30 %. Feed size was 1.5 mm for the first 20 days and 2 mm for the remaining feeding period.

At the middle (Day 31) and end of the experiment (Day 55) whiteleg shrimp from each tank were weighed to determine mean body weight, relative growth rate (RGR), feed intake, feed conversion ratio (FCR) and survival. Additionally, 5 shrimp per tank were sampled for whole-body composition analysis at the end of the experiment. At the indicated times, shrimps were sorted and euthanized for tissue sampling. From three individuals per tank (15 per experimental group), fresh haemolymph was collected for haemocytes total counting and plasma for the assessment on innate immune parameters. Hepatopancreas was also collected for analysis of oxidative status and gene expression analysis of key immune related genes through RT-qPCR (Table 3).

**Table 3.** Immune-related genes analysed by real-time quantitative PCR and biological function.

| Gene | Biological function |
| --- | --- |
| Cytoplasmic-type actin 4 | Cell motility, ubiquitously expressed in all eukaryotic (house-keeping gene) |
| Penaeidin-3a | Antibacterial activity |
| C-type lectin 2-like | Cell-cell adhesion, immune response to pathogens and apoptosis. |
| Hemocyanin | Oxygen transport |
| Rab GTPase | Formation, transport and fusion of intracellular organelles key for immune response |
| Lysozyme C-like | Bacteriolytic function |
| Glutathione transferase | Antioxidant defence (xenobiotics detoxification) |
| Glutathione peroxidase | Antioxidant defence (organic and inorganic peroxides) |
| Crustin p | Peptidase inhibitor activity |
| Thioredoxin 1 | Redox signalling |
| Caspase 3 | Apoptosis signalling (executioner caspase) |
| Spermine synthase | Cell cycle and proliferation (polyamine biosynthesis) |
| Adenosylhomocysteine hydrolase | Intracellular regulation of homocysteine formation |

With the aim of assessing the modulatory roles of specific phytochemicals in the inflammatory processes, the inflammatory response was further studied at the end the 55 days feeding period. Twenty-five shrimps were re-allocated in a similar recirculation system where the diets were established in triplicate tanks and the individuals were submitted to a challenge bath in 20 L tanks with *Vibrio parahaemolyticus* (2 x 10^6^ CFU mL^-1^) for 1 h. After that, food was replenished according to the previous regime and mortality recorded for one week. Moreover, 5 individuals from each tank (15 per group) were sampled for haemolymph and hepatopancreas at 4, 24 and 48 hours after the challenge period in order to assess cell proliferation and plasma immune mechanisms.

### 2.3. Bromatological analysis

Bromatological analysis of the experimental diets was performed following the Association of Official Analytical Chemists procedures ^18^. Briefly, dry matter was determined by drying samples at 105 °C in an oven until constant weight; ash, by incineration at 450 °C for 16 h in a muffle furnace; crude protein content (N x 6.25), by the Kjeldahl method after acid digestion using a Kjeltec digestion and distillation unit; lipid content, by petroleum ether extraction (Soxtec HT System) and gross energy, by direct combustion in an adiabatic bomb calorimeter (PARR Instruments, Moline, IL, USA; PARR model 1261).

### 2.4. Chemical profile of shrimp carcasses

#### 2.4.1. Pre-treatment of the samples

The samples were stored at -80 °C until starting the pre-treatment of the samples. The pre-treatment consisted of separating the muscle from the exoskeleton. All animals (shrimp) belonging to the same group were together. Then the muscle was cut into very small pieces and placed in an oven at 35 °C, to dry/dehydrate, until the weight of the samples stabilized. Finally, the samples taken from the oven were macerated, using a mill, and stored.

#### 2.4.2. Extracts preparation

Nearly 2.5 g of dried biomass of the sample were extracted with hexane (in a proportion of 1 g : 15 mL), at room temperature with stirring for 72 h. After the first extraction, the solvent is renewed and the extraction continues for another 72 h. The solvent from the combined hexane extractions was evaporated to dryness ^19^.

#### 2.4.3. Trimethylsilyl derivatives preparation

Before analysis by gc-ms, samples were derivatized according to the known methodology and usually used in our department with slight changes ^20^. To 10 mg of extract, 125 μL of pyridine, 125 μL of N, O-bis(trimethylsilyl)trifluoroacetamide (BSTFA), and 25 μL of trimethylchlorosilane (TMSCl) were added. 200 μL of 1 mg mL-1 of hexatriacontane (internal standard) and 525 μL of dichloromethane were added, making a total volume of 1mL. The mixtures were maintained at 70 ℃ for 30 minutes and then were transferred to vial, with needles and syringes. The samples were injected into the GC-MS apparatus. All extracts were derivatized in triplicate and injected in duplicate.

#### 2.4.4. Gas Chromatography-Mass Spectrometry Analysis

GC–MS analysis of each silylated sample was performed using a GC–MS QP2010 Ultra Shimadzu (University of Aveiro, Portugal) equipped with a ZB-5ms J & W capillary column (30 m x 0.25 mm x 0.25 μm) as described in Ferreira, et al. ^21^. Samples were injected with a split ratio of 1:10 and helium as carrier gas with a flux of 1.19 mL/min. The injector temperature was at 320 °C and the transfer-line temperature was at 200 °C. The temperature of the column was maintained at 70 °C for 5 mins and then increased, first at 4 °C min^-1^. until 250 °C, followed by an increase of 2 °C min^-1^. until 300 °C, which was maintained for 5 min. The mass spectrometer was operated in the electronic impact (EI) mode with an energy of 0.1 kV, and data were collected at a rate of 1scan/s over a range of m/z 50–1000. The performed chromatography lasted 80 min and the standards were analyzed separately by GC–MS under the same chromatographic conditions. The compounds were identified based on a direct comparison with the mass spectra database libraries (NIST14 Mass Spectra and WILEY Registry TM of Mass Spectra Data)

### 2.5. Tissue collection

Haemolymph was extracted from the ventral part of the first abdominal segment with an insulin syringe containing 200 µL of anticoagulant solution (27 mM Trisodium citrate, 385 mM sodium chloride, 115 mM glucose in mQ water, pH 7.5).

Immediately after extraction, haemolymph was diluted in anticoagulant solution 1:1 (v/v). 20 µL, 50 µL and 200 µL of haemolymph were used for oxyhemocyanin determination, total haemocyte counting and respiratory burst analysis respectively. The remaining haemolymph was then centrifuged at 2500 rpm, 10 min at 4 °C and plasma was transferred to new 1.5 mL Eppendorf and freeze immediately in liquid nitrogen.

Hepatopancreas was collected and cut in two longitudinal pieces, then one piece was placed in 1.5 mL tube with 1 mL of RNA later (Sigma). Samples were then maintained 24 hours at 4 °C and then at -20 °C until RNA extraction.

For the oxidant status assessment, shrimp hepatopancreas was collected, immediately frozen in liquid nitrogen, and then stored in -80 °C until further analyses.

### 2.6. Haemocytes count (THC)

Briefly, 50 µL of 1:1 diluted haemolymph with anticoagulant solution were transferred to a new 1.5 mL tube with 50 µL of formalin according to Bautista-Covarrubias, et al. ^22^ to a final dilution of 1:4. Samples were counted in a Neubauer chamber under a light microscope at 400x.

### 2.7. Haemolymph humoral parameters

#### 2.7.1. Oxyhemocyanin

For hemocyanin measurements, 20 µL hemolymph was immediately diluted with 180 µL distilled water. Then, 100 µL were transferred to a microplate by duplicate and read at 335 nm. Hemocyanin concentration was calculated using an extinction coefficient of 17.26 calculated on the basis of the 74000 Da functional subunit.

#### 2.7.2. Respiratory burst (RB)

Intracellular production of the super oxide anion in haemocytes was quantified using the NBT (Nitroblue Tetrazolium) reduction to formazan was measured according to the Fujiki and Yano ^23^ method of with some adaptations. Briefly, 100 µL of hemolymph diluted in 500 µL of cacodilate buffer solution (10 mM cacodylate, 10 mM CaCl_2_, pH 7.0) was placed in a well of microplate (for triplicate) and centrifuged at 800 × g for 10 min. Plasma was removed and hemocytes washed with 100 μL Hank’s solution. Next, 100 μL of *Vibrio anguillarum* inactivated bacteria at 10^6^ cells per mL was added and incubated for 2 h at room temperature. The bacteria solution was then removed and haemocytes washed three times with 100 μL Hank’s solution and stained for 30 min with NBT solution (0.3 %) at room temperature. After fixed, haemocytes were washed three times with 100 μL methanol (70 %) and dried for 5 min. Formazan was dissolved with 120 μL KOH and 140 μL DMSO, and absorbance was read at 630 nm using a Synergy HT microplate reader (Biotek).

### 2.8. Plasma immune parameters

Hemolymph which was not used for respiratory burst, oxyhemocyanin determination and THC was centrifuged at 2500 × rpm for 10 min at 4 °C. Plasma was placed in a new tube and hemocytes were discarded. Two µL of plasma were used to measure protein concentration in the DeNovix DS-11 spectrophotometer according to manufacturer recommendations.

Briefly, pro-phenoloxidase activity was measured as follows: 25 μL of plasma were incubated for 30 min at 25 °C with 100 μL trypsin (1 mg mL^-1^). Then, 100 μL L-DOPA (3 mg mL^-1^) were added and absorbance was measured each minute during 5 minutes at 490 nm in a Synergy HT microplate reader. Results were calculated using the Beer-Lambert law by means of the molar extinction coefficient of the L-DOPA (3700) according to Ji, et al. ^24^. Results are expressed as units of pro-phenoloxidase per mL of plasma.

Lysozyme activity was measured using a turbidimetric assay according to Costas, et al. ^25^. Briefly, a solution of *Micrococcus lysodeikticus* (0.25 mg mL^-1^, 0.05 M sodium phosphate buffer, pH 6.2) was prepared. To a microplate, 20 mL of plasma in duplicates and 170 µL of the above suspension were added to give a final volume of 190 µL. The reaction was carried out at 25 °C and the absorbance (450 nm) was measured after 0 and 10 min in a Synergy HT microplate reader. Lyophilized hen egg white lysozyme (Sigma-Aldrich) was serially diluted in sodium phosphate buffer (0.05 M, pH 6.2) and used to develop a standard curve. The amount of lysozyme in the sample was calculated using the formula of the standard curve.

The nitric oxide production was assayed in plasma by the Griess reaction. This reaction quantifies the nitrite content of the sample of 25 µL of plasma by adding 100 µL of 1 % sulfanilamide (Sigma-Aldrich) in 2.5 % of phosphoric acid to each well, followed by 100 µL of 0.1 % N-naphthyl-ethylenediamine (Sigma) in 2.5 % phosphoric acid. After 10 min of incubation at room temperature, the optical density was determined in a Synergy HT microplate reader (Biotek) at 540 nm. The concentration of nitrite in the sample was determined from standard curves generated using known concentrations of sodium nitrite.

Antiprotease activity was quantified using the azocasein hydrolysis assay according to Ramos-Pinto, et al. ^26^ with some modifications. Briefly, 20 μl of plasma were added to a microtube with 10 µL of trypsin (5 mg mL^-1^). After 10 minutes incubation, 90 µL of PBS and 125 µL of azocasein at 2 % in a 100 mM ammonium bicarbonate buffer were added and incubate for 1 hour at 25 °C. The reaction was stopped by adding 250 µL of 10 % trichloro acetic acid (TCA) and the mixture was centrifuged (10,000 × g, 5 min). Then, 100 µL of the supernatant solution were transferred to a 96-well plate in duplicate containing 100 μL well^−1^ of 1N NaOH. The optical density (OD) was read at 450 nm using a Synergy HT microplate reader (Biotek). Plasma was replaced by PBS as reference sample (100 % of antiprotease activity), and sample with 20 µL of PBS was used as blank (0 % activity). Results are expressed as the ability to suppress the trypsin activity against the reference sample.

Protease activity was quantified using the azocasein hydrolysis assay according to Ramos-Pinto, et al. ^26^ with some modifications. Briefly, 20 μl of plasma were incubated with 90 µL of PBS and 125 µL of azocasein at 2 % in a 100 mM ammonium bicarbonate buffer for 24 hours at 24 °C in microtubes with continuous shaking, protected from direct light. The reaction was stopped by adding 250 µL of 10 % TCA and the mixture was centrifuged (6.000 × g, 5 min). Then, 100 µL of the supernatant solution were transferred to a 96-well plate in duplicate containing 100 μL well^−1^ of 1N NaOH. The OD was read at 450 nm using a Synergy HT microplate reader (Biotek). Plasma was replaced by PBS as reference sample (100 % of protease activity), and sample with 20 µL of PBS was used as blank (0 % activity). Results are expressed as the ability to suppress the trypsin activity against the reference sample.

### 2.9. Hepatopancreas oxidant status

All measurements were carried out in a SpectraMax 190 microplate reader, at 25 °C. Shrimp hepatopancreas were homogenised using a Potter–Elvehjem homogenizer, in chilled phosphate buffer (0.1 M, pH 7.4) in a 1:11 ratio [tissue mass (mg): buffer volume (mL)]. The resulting homogenate was then divided into two aliquots, *i.e*. for lipid peroxidation (LPO) measurement and post-mitochondrial supernatant (PMS) preparation. The PMS was obtained by centrifugation in a refrigerated centrifuge (Eppendorf 5415R) at 13,400 × g for 20 min at 4 °C. Aliquots of PMS were then divided into microtubes and stored at -80 °C until further analyses. Superoxide dismutase (SOD), catalase (CAT), and glutathione peroxidase (GPx) activities as well as lipid peroxidation (LPO) levels were measured.

SOD activity was assayed in PMS with a Ransod kit (Randox Laboratories Ltd., UK). The method employs xanthine and xanthine oxidase to generate superoxide radicals, which react with 2-(4-iodo-phenyl)-3-(4-nitrophenol)-5-phenyltetrazolium chloride (INT) to form a red formazan dye determined at 505 nm. SOD activity was then measured by the degree of inhibition of this reaction, considering that one unit of SOD causes a 50 % inhibition of the rate of reduction of INT, under the conditions of the assay. Results were expressed as SOD unit’s mg protein^-1^.

CAT activity was assayed in PMS by the method of ^27^, with slight modifications. Briefly, the assay mixture consisted of 0.190 mL phosphate buffer (0.05 M, pH 7.0) with hydrogen peroxide (H_2_O_2_; 0.010 M) and 0.010 mL of PMS, in a final volume of 0.2 mL. Change in absorbance was measured in appropriated UV transparent microplates (UV-Star® flat-bottom microplates, Greiner Bio-One GmbH, Germany), recorded at 240 nm and CAT activity was calculated in terms of μmol H_2_O_2_ consumed min^-1^ mg^-1^ protein using a molar extinction coefficient (ε) of 43.5 M^-1^ cm^-1^.

GPx activity was determined in PMS according to the method described by Mohandas, et al. ^28^ and modified by Athar and Iqbal ^29^. The assay mixture consisted of 0.09 mL phosphate buffer (0.05 M, pH 7.0), 0.03 mL ethylenediaminetetraacetic acid (EDTA; 0.010 M), 0.03 mL sodium azide (0.010 M), 0.03 mL glutathione reductase (2.4 U mL^-1^), 0.03 mL reduced glutathione (GSH; 0.010 M), 0.03 mL nicotinamide adenine dinucleotide phosphate-oxidase (NADPH; 0.0015 M), 0.03 mL H_2_O_2_ (0.0025 M) and 0.03 mL of PMS in a total volume of 0.3 mL. Oxidation of NADPH to NADP^+^ was recorded at 340 nm in a SpectraMax 190 microplate reader and GPx activity was calculated in terms of nmol NADPH oxidized min^-1^ mg protein^-1^ using a ε of 6.22 × 10^3^ M^-1^ cm^-1^.

As an estimation of LPO, TBARS quantification was carried out in the previously prepared homogenate according to the procedure of ^30^ and adapted by ^31,32^ and Aloísio Torres, et al. ^33^. Briefly, 0.005 mL of butylatedhydroxytoluene (BHT; 4% in methanol) and 0.045 mL of phosphate buffer (0.05 M, pH 7.4) were added to 0.05 mL of homogenate and mixed well to prevent oxidation. To this aliquot, 0.250 mL of TCA (12%), 0.225 mL of Tris–HCl (0.060 M, pH 7.4 and 0.0001 M DTPA) and 0.250 mL of thiobarbituric acid (TBA; 0.73 %) were added and well mixed. This mixture was heated for 1 h in a water bath set at 100°C and then cooled to room temperature, and centrifuged at 15,700 × g for 5 min. The absorbance of each sample was measured at 535 nm in a SpectraMax 190 microplate reader. The rate of LPO was expressed in nmol of thiobarbituric acid reactive substances (TBARS) formed mg protein^-1^ using a ε of 1.56 × 10^5^ M^-1^ cm^-1^.

### 2.10. Hepatopancreas gene expression

In a first approach, cDNA was only isolated from the hepatopancreas of shrimps sampled at 31 and 55 days fed the experimental diets. This procedure followed Machado, et al. ^34^ with some modifications. Briefly, hepatopancreas total RNA isolation and DNase treatment (NZY Total RNA isolation kit, MB13402, NZYTech, Portugal) and first-strand cDNA synthesis (NZY First-strand cDNA synthesis kit, MB125, NZYTech, Portugal) were performed according to manufacture guidelines. Primers design and efficiency values and quantitative PCR assays were performed. Efficiency of each primer was calculated according to a series dilution of hepatopancreas cDNA pool. Primers efficiency and DNA amplification was carried out using the CFX384 Touch Real-Time PCR Detection System with specific primers for genes that have been selected for their involvement in immune responses (Table 3). Accession number, efficiency values, annealing temperature, product length and primers sequences are presented in Table 4. Melting curve analysis was also performed to verify that no primer dimers were amplified. The standard cycling conditions were 95 °C initial denaturation for 10 min, followed by 40 cycles of 94 °C denaturation for 30 s, primer annealing temperature for 30 s and 72 °C extension for 30 s. All reactions were carried out as technical duplicates. The expression of the target genes was normalised using the expression of whiteleg shrimp cytoplasmic-type actin 4 (actin).

**Table 4.**
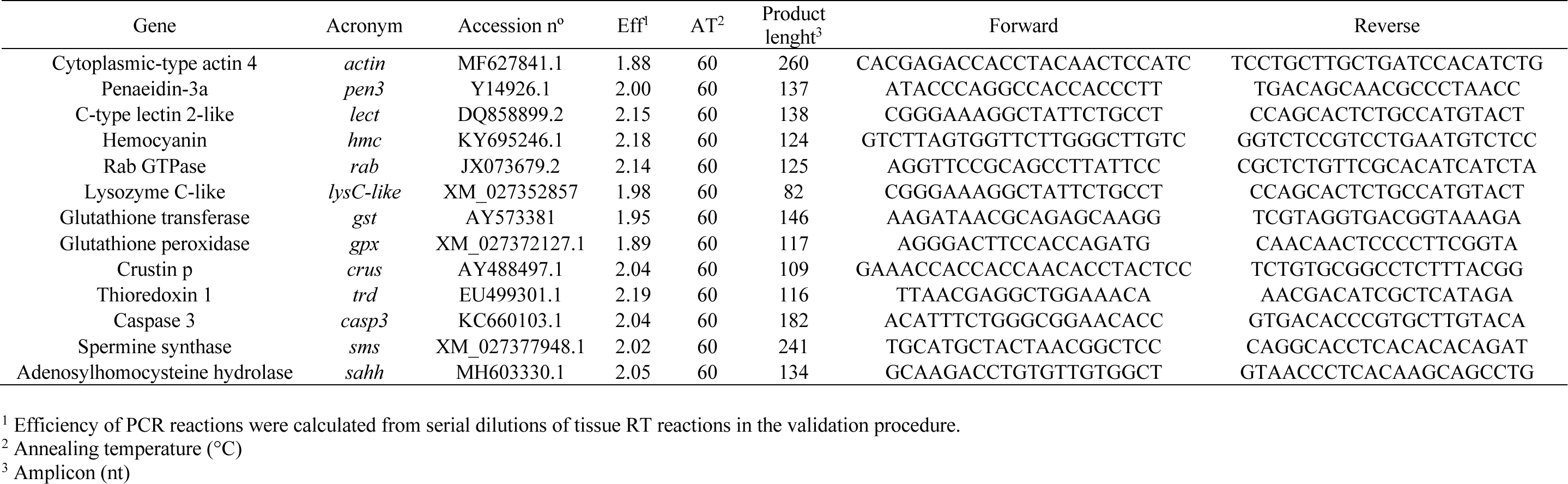
Forward and reverse primers for real-time quantitative PCR.

### 2.11. Bacterial challenge

For the bacterial challenge, *Vibrio parahaemolyticus*, kindly provided by Prof. Miguel Ángel Moríñigo (University of Malaga), was used. Bacteria was routinely cultured at 25 °C in tryptic soy broth (TSB) or tryptic soy agar (TSA) supplemented with NaCl to a final concentration of 2 % (w/v) and stored at -80 °C in TSB 2 % NaCl with 15 % (v/v) glycerol. To prepare the inoculum for the challenge bath, stock bacteria was cultured overnight at 25 °C in TSB 2 % NaCl and exponentially growing bacteria were collected and adjusted to a final concentration of 2 × 10^8^ colony forming units (cfu) mL^-1^, as confirmed by plating the resulting culture on TSA – 2 % NaCl plates and counting of cfu. The recirculation water system was stopped, and the water volume of each tank was lowered to a final volume of 10 L. Shrimp were inoculated with the bacteria with strong aeration for 1 h. Afterwards, the rearing water was changed three times and the recirculation system was re-established.

### 2.12. Data analysis

#### 2.12.1. Growth performance

Relative growth rate (RGR, % weight day^-1^) was calculated as: RGR = (eg^–1^) × 100, where g = (lnWf – lnWi) × t^–1^. Wf and Wi correspond to the final and initial weights, respectively. Feed conversion ratio (FCR) was calculated as: FCR = (Fi / Wg), where Fi corresponds to feed intake (g) and Wg to the mean weight gain (g). Survival was expressed as percentage and calculated as: S = (Sf / Si) × 100, where Si and Sf correspond to the initial and final number of individuals in the tanks, respectively. Differences in growth performance, survival, whole-body composition, between dietary treatments were evaluated using One-way ANOVA’s, followed by Tukey HSD multiple comparison tests. Kruskal-Wallis one-way analysis of variance tests, followed by Wilcoxon pairwise comparison tests were used when data did not comply with the One-way ANOVA’s assumptions.

#### 2.12.2. Humoral, oxidative stress and chemical profile

Differences in oxidative stress biomarkers, immune parameters and gene expression and chemical profile between dietary treatments were evaluated using One-way ANOVA’s, followed by Tukey HSD multiple comparison tests. Kruskal-Wallis one-way analysis of variance tests, followed by Wilcoxon pairwise comparison tests were used when data did not comply with the One-way ANOVA’s assumptions.

To evaluate the activation of the inflammatory mechanisms, the sampling point 55 days was used as time 0h during the time-course data analysis, as they represent unstimulated animals prior to infection. Differences between dietary treatments during the time course period were evaluated using Two-way ANOVA’s, followed by Tukey HSD multiple comparison tests. In an attempt to discriminate and classify the dietary treatments along the experimental sampling points, a multivariate canonical discriminant analysis using XLSTAT (Addinsoft, New York, USA), was performed on the dataset to evaluate the linear combinations of the original variables that will best separate the groups (discriminant functions). Multivariate linear discriminant analyses (LDA) were performed for gene expression data to evaluate how it contributed to the dissociation of the diets in the discriminant functions generated. A MANOVA was performed to assess discriminatory significance using Wilk’s λ test, after checking data compliance to the statistics assumptions. The distance between group centroids was measured by Mahalanobis distance and its significance inferred by One-way ANOVA’s statistics. Results were expressed as means ± standard deviation (SD). In results expressed as percentage, an arcsine transformation was performed prior to any statistical test: T = ASIN (SQRT (value / 100)). The significance level considered was P < 0.05 for all tests performed.

## 3. Results

### 3.1. Feeding trial

#### 3.1.1. Growth performance

At day 31 of feeding, no significant differences between treatments were found in final body weight, RGR and survival values. However, whiteleg shrimps fed SS10 and SL10 diets had a significantly higher FCR values than those fed the CTRL diet, while shrimp fed SL10 had significantly higher FCR than those fed the remaining diets, excluding SS10. Feed intake was also higher in whiteleg shrimp fed the experimental diets rich in *S. ramosissima* biomasses than those fed CTRL, particularly shrimp fed diet SL10 had significantly higher feed intake values than those fed the remaining diets, excluding SS10 (Table 5).

**Table 5.** Initial and final weight, relative growth rate (RGR), feed conversion ratio (FCR), feed intake and survival of juvenile whiteleg shrimp (*P. vannamei*) fed the experimental diets for 31 days.

|  | CTRL | SS5 | SS10 | SL5 | SL10 |
| --- | --- | --- | --- | --- | --- |
| Initial weight (g) | 6.1 ± 0.0 | 6.1 ± 0.1 | 6.0 ± 0.0 | 6.1 ± 0.0 | 6.1 ± 0.1 |
| Final weight (g) | 12.9 ± 0.3 | 13.2 ± 0.3 | 12.9 ± 0.2 | 13.2 ± 0.3 | 13.1 ± 0.2 |
| RGR (% day<sup>-1</sup>) | 2.4 ± 0.1 | 2.5 ± 0.1 | 2.5 ± 0.0 | 2.5 ± 0.1 | 2.5 ± 0.0 |
| FCR | 2.5 ± 0.1<sup>a</sup> | 2.7 ± 0.1<sup>ab</sup> | 2.9 ± 0.1<sup>bc</sup> | 2.7 ± 0.1<sup>ab</sup> | 3.0 ± 0.1<sup>c</sup> |
| Feed intake (% ABW day<sup>-1</sup>) | 5.7 ± 0.3<sup>a</sup> | 6.3 ± 0.1<sup>b</sup> | 6.7 ± 0.2<sup>bc</sup> | 6.5 ± 0.2<sup>b</sup> | 7.0 ± 0.3<sup>c</sup> |
| Survival (%) | 100.0 ± 0.0 | 99.6 ± 0.7 | 99.6 ± 0.7 | 98.9 ± 1.5 | 99.6 ± 0.7 |

Results expressed as mean ± standard deviation. For initial weight n = 55 and for the remaining parameters n = 5 experimental units. Different superscript letters indicate statistical differences ( $P < 0.05$ ) between treatments in a One-way ANOVA, followed by a Tukey HSD multiple comparison test.

Moreover, no significant differences between treatments were observed in final body weight, RGR and survival values after 31 days of feeding. Still, whiteleg shrimps fed the CTRL diet had significantly lower FCR and feed intake values than the experimental diets where *S. ramosissima* biomasses were included, while shrimp fed SL10 had significantly higher FCR and feed intake values than those fed the remaining diets (Table 6).

**Table 6.** Initial and final weight, relative growth rate (RGR), feed conversion ratio (FCR), feed intake and survival of juvenile whiteleg shrimp (*P. vannamei*) fed the experimental diets for 55 days.

|  | CTRL | SS5 | SS10 | SL5 | SL10 |
| --- | --- | --- | --- | --- | --- |
| Initial weight (g) | 6.1 ± 0.0 | 6.1 ± 0.1 | 6.0 ± 0.0 | 6.1 ± 0.0 | 6.1 ± 0.1 |
| Final weight (g) | 17.6 ± 0.5 | 18.1 ± 0.5 | 18.1 ± 0.4 | 18.1 ± 0.4 | 18.3 ± 0.3 |
| RGR (% day<sup>-1</sup>) | 2.0 ± 0.0 | 2.0 ± 0.1 | 2.0 ± 0.0 | 2.0 ± 0.0 | 2.0 ± 0.0 |
| FCR | 3.1 ± 0.2<sup>a</sup> | 3.6 ± 0.2<sup>b</sup> | 3.7 ± 0.1<sup>b</sup> | 3.6 ± 0.2<sup>b</sup> | 4.0 ± 0.1<sup>c</sup> |
| Feed intake (% ABW day<sup>-1</sup>) | 5.3 ± 0.3<sup>a</sup> | 6.2 ± 0.2<sup>b</sup> | 6.4 ± 0.2<sup>b</sup> | 6.3 ± 0.2<sup>b</sup> | 7.1 ± 0.2<sup>c</sup> |
| Survival (%) | 96.7 ± 1.4 | 97.1 ± 2.2 | 98.2 ± 2.3 | 96.0 ± 2.4 | 97.8 ± 1.4 |

Results expressed as mean ± standard deviation. For initial weight n = 55 and for the remaining parameters n = 5 experimental units. Different superscript letters indicate statistical differences ( $P < 0.05$ ) between treatments in a One-way ANOVA, followed by a Tukey HSD multiple comparison test.

At the end of the feeding trial (Day 55), significant differences between treatments were found in the shrimp whole-body composition. Shrimp fed the CTRL and SL5 diets had significantly higher dry matter contents than those fed SL10, while those fed SS10 had significantly higher ash contents than SL5 (Table 7).

**Table 7:** Whole-body composition (% fresh weight) of juvenile whiteleg shrimp (*P. vannamei*) fed the experimental diets for 55 days.

|  | CTRL | SS5 | SS10 | SL5 | SL10 |
| --- | --- | --- | --- | --- | --- |
| Dry matter (%) | 24.6 ± 1.1<sup>b</sup> | 23.3 ± 0.4<sup>ab</sup> | 23.4 ± 0.6<sup>ab</sup> | 24.3 ± 0.5<sup>b</sup> | 22.9 ± 0.7<sup>a</sup> |
| Protein (%) | 16.8 ± 1.2 | 17.1 ± 0.3 | 17.2 ± 0.4 | 17.5 ± 0.6 | 16.8 ± 0.6 |
| Lipids (%) | 1.7 ± 0.4 | 1.5 ± 0.2 | 1.3 ± 0.1 | 1.6 ± 0.2 | 1.3 ± 0.3 |
| Ash (%) | 4.0 ± 0.3<sup>ab</sup> | 4.0 ± 0.6<sup>ab</sup> | 5.3 ± 1.2<sup>b</sup> | 3.7 ± 0.2<sup>a</sup> | 4.7 ± 0.5<sup>ab</sup> |
| Energy (kJ g<sup>-1</sup>) | 5.0 ± 0.2 | 4.9 ± 0.1 | 4.8 ± 0.2 | 5.0 ± 0.2 | 4.7 ± 0.2 |

Results expressed as mean ± standard deviation (n = 5 experimental units). Different superscript letters indicate statistical differences ( $P < 0.05$ ) between treatments in a One-way ANOVA, followed by a Tukey HSD multiple comparison test.

#### 3.1.2. Phytochemical profile

The phytochemical characterization of whiteleg shrimp allowed the identification of 50 and 40 compounds, after 31 days (Table 8) and 55 days of feeding (Table 9), respectively. The identified compounds mainly belong to 5 classes, namely alcohols, amino acids, fatty acids, organic acids and sterols. From the analyses of the phytochemical profile is possible to highlight that at 31 days of feeding, the total relative abundance of both alcohols and sterols is lower in the groups fed with stems and leaves and seeds than in the CTRL group. In the case of alcohols, in groups SS10, SS5 and SL5, this difference is statistically significant. For sterols, only the SL5 and SL10 groups are significantly lower than the CTRL. In organic acids and fatty acids there are significant differences between CTRL and the SL10 group: the relative abundance of organic acids is higher in the SL10 group than in the CTRL, while the relative abundance of fatty acids is lower in the SL10 than in the CTRL. Finally, the amino acids in the SS5, SL5, and SL10 groups have significantly higher relative abundances than CTRL. Regarding the phytochemical profile at 55 days of feeding, the total relative abundance of alcohols is significantly greater in the SS10 and SL5 groups than in the CTRL. In the case of amino acids, the SL10 group has a statistically greater abundance than in CTRL. The SL5 group, draws attention, having a total relative abundance of amino acids much lower than CTRL and even in relation to the other groups. When looking at this particular situation it appears that there is an abrupt reduction in the relative abundance of glycine in this group.

**Table 8:** Compounds identification from the hexane extracts of whiteleg shrimp (*P. vannamei*) fed dietary treatments after 31 days. Values are presented in relative abundance (%) (means ± SD (n = 6)). P-values from Kruskal-Wallis (p ≤ 0.05).

| Identified compounds |  | Dietary treatments |  |  |  |  |
| --- | --- | --- | --- | --- | --- | --- |
|  |  | 31 Days |  |  |  |  |
|  |  | CTRL | SS5 | SS10 | SL5 | SL10 |
| **Alcohol** |  |  |  |  |  |  |
|  | 1-Hexadecanol (%) | 0.3 ± 0.05 | 0.09 ± 0.01 | 0.05 ± 0.05 | 0.21 ± 0.03 | 0.27 ± 0.02 |
|  | 1-Octadecanol (%) | 0.19 ± 0.01 | 0.03 ± 0.03 | 0.02 ± 0.03 | 0.12 ± 0.03 | 0.16 ± 0.02 |
|  | 9-Octadecen-1-ol (%) | 0.27 ± 0.05 | 0.11 ± 0.02 | 0.12 ± 0 | 0.16 ± 0.03 | 0.22 ± 0.04 |
|  | gamma-Tocopherol (%) | 0.41 ± 0.01 | 0.33 ± 0.03 | 0.36 ± 0.03 | 0.25 ± 0.01 | 0.27 ± 0.01 |
|  | Glycerol (%) | 0.53 ± 0.11 | 0.52 ± 0.03 | 0.45 ± 0.21 | 0.64 ± 0.15 | 0.62 ± 0.16 |
| **Total** |  | **1.69 ± 0.04** | **1.08 ± 0.02** | **1 ± 0.06** | **1.38 ± 0.05** | **1.53 ± 0.05** |
| **Amino acid** |  |  |  |  |  |  |
|  | Alanine (%) | 1.2 ± 0.01 | 2.53 ± 0.15 | 1.27 ± 0.08 | 4.01 ± 1.08 | 2.93 ± 0.61 |
|  | Asparagine (%) | 0.11 ± 0.01 | 0.22 ± 0.04 | 0.11 ± 0.05 | 0.31 ± 0.08 | 0.19 ± 0.07 |
|  | Glycine (%) | 3.79 ± 0.45 | 4.63 ± 0.35 | 4.09 ± 1.2 | 6.03 ± 2.49 | 7.97 ± 2.32 |
|  | Hydroxyproline (%) | 0.08 ± 0.01 | 0.11 ± 0 | 0.09 ± 0 | 0.13 ± 0.03 | 0.19 ± 0.09 |
|  | Leucine (%) | 0.11 ± 0 | 0.39 ± 0.22 | 0.13 ± 0.05 | 0.66 ± 0.29 | 0.25 ± 0.06 |
|  | Lysine (%) | 0.43 ± 0.06 | 0.66 ± 0.26 | 0.49 ± 0.07 | 1.02 ± 0.23 | 0.88 ± 0.14 |
|  | Ornithine (%) | 0.75 ± 0.05 | 0.91 ± 0.08 | 0.95 ± 0.12 | 1.43 ± 0.37 | 1.5 ± 0.16 |
|  | Phenylalanine (%) | 0.38 ± 0.13 | 1.35 ± 0.3 | 0.37 ± 0.14 | 1.54 ± 0.38 | 1.11 ± 0.15 |
|  | Proline (%) | 2.68 ± 0.1 | 5.03 ± 0.09 | 0.82 ± 0.53 | 0.94 ± 0.46 | 5.42 ± 0.8 |
|  | Serine (%) | 0.05 ± 0 | 0.16 ± 0.04 | 0.03 ± 0 | 0.17 ± 0.05 | 0.12 ± 0.03 |
|  | Threonine (%) | 0.23 ± 0.01 | 0.59 ± 0.09 | 0.2 ± 0.04 | 0.67 ± 0.21 | 0.52 ± 0.13 |
|  | Tyrosine (%) | 0.19 ± 0.02 | 0.39 ± 0.13 | 0.12 ± 0.02 | 0.21 ± 0.06 | 0.25 ± 0.02 |
|  | Valine (%) | 0.55 ± 0.01 | 2.07 ± 0.71 | 0.61 ± 0.05 | 3.84 ± 1 | 1.87 ± 0.05 |
| **Total** |  | **10.55 ± 0.07** | **19.03 ± 0.19** | **9.28 ± 0.18** | **20.97 ± 0.52** | **23.18 ± 0.36** |
| **Fatty acid** |  |  |  |  |  |  |
|  | 11.14-Eicosadienoic acid (%) | 1.32 ± 0.09 | 1.3 ± 0.14 | 1.14 ± 0.15 | 1.48 ± 0.51 | 0 ± 0 |
|  | 11-Eicosaenoic acid (%) | 0.54 ± 0.03 | 0.5 ± 0.02 | 0.59 ± 0.03 | 0.58 ± 0.09 | 0.43 ± 0.05 |
|  | Arachidonic acid (%) | 0.75 ± 0.01 | 0.62 ± 0.06 | 0.78 ± 0.03 | 0.69 ± 0.06 | 0.62 ± 0.03 |
|  | Doconexent (DHA) (%) | 3.86 ± 0.05 | 3.26 ± 0.21 | 3.98 ± 0.27 | 3.92 ± 0.34 | 3.23 ± 0.31 |
|  | Eicosapentaenoic Acid (EPA) (%) | 4.43 ± 0.03 | 3.96 ± 0.27 | 4.72 ± 0.3 | 5.29 ± 0.56 | 4.09 ± 0.44 |
|  | Heptadecanoic acid (%) | 0.47 ± 0.01 | 0.45 ± 0 | 0.52 ± 0 | 0.5 ± 0.02 | 0.38 ± 0.02 |
|  | Linoleic acid (%) | 9.14 ± 0.21 | 8.65 ± 0.26 | 10 ± 0.66 | 9.64 ± 1.22 | 8.28 ± 0.85 |
|  | Nonadecanoic acid (%) | 0.16 ± 0.03 | 0.17 ± 0.03 | 0.25 ± 0.02 | 0.17 ± 0.03 | 0.16 ± 0.03 |
|  | Oleic Acid (%) | 9.32 ± 0.5 | 9.32 ± 1.19 | 9.79 ± 0.4 | 9.88 ± 1.87 | 7.99 ± 1.13 |
|  | Palmitelaidic acid (%) | 0.36 ± 0.02 | 0.34 ± 0.05 | 0.4 ± 0.03 | 0.41 ± 0.05 | 0.38 ± 0.03 |
|  | Palmitic Acid (%) | 9.27 ± 0.25 | 8.76 ± 0.34 | 9.83 ± 0.2 | 9.89 ± 0.55 | 8.07 ± 0.73 |
|  | Pentadecanoic acid (%) | 0.1 ± 0 | 0 ± 0 | 0.12 ± 0 | 0.12 ± 0.01 | 0.12 ± 0.02 |
|  | Stearic acid (%) | 5.59 ± 0.2 | 5.1 ± 0.21 | 5.68 ± 0.41 | 5.45 ± 0.09 | 4.53 ± 0.4 |
| **Total** |  | **45.32 ± 0.11** | **42.44 ± 0.21** | **47.79 ± 0.19** | **48.01 ± 0.41** | **38.28 ± 0.31** |
| **Organic acid** |  |  |  |  |  |  |
|  | 2-Hydroxyisocaproic acid (%) | 0.16 ± 0.02 | 0.22 ± 0.02 | 0.2 ± 0.03 | 0.26 ± 0.08 | 0.23 ± 0.07 |
|  | Aminomalonic acid (%) | 0.12 ± 0.03 | 0.13 ± 0.07 | 0.17 ± 0.04 | 0.24 ± 0.14 | 0.22 ± 0.11 |
|  | Behenic acid (%) | 0.12 ± 0.01 | 0.14 ± 0 | 0.12 ± 0 | 0.13 ± 0.03 | 0.12 ± 0 |
|  | Glyceric acid (%) | 0.06 ± 0.01 | 0.09 ± 0.01 | 0.06 ± 0.01 | 0.12 ± 0.03 | 0.1 ± 0.03 |
|  | Lactic Acid (%) | 1.31 ± 0.22 | 1.53 ± 0.1 | 1.25 ± 0.33 | 1.35 ± 0.52 | 2 ± 0.76 |
|  | Malic acid (%) | 0.11 ± 0.01 | 0.12 ± 0.02 | 0.08 ± 0.01 | 0.22 ± 0.06 | 0.14 ± 0.05 |
|  | Pentanedioic acid (%) | 1.17 ± 0.1 | 1.42 ± 0.12 | 1.03 ± 0.28 | 1.33 ± 0.45 | 1.67 ± 0.54 |
|  | Phosphoric acid (%) | 0.12 ± 0.02 | 0.26 ± 0.04 | 0.17 ± 0.01 | 0.39 ± 0.11 | 0.27 ± 0.07 |
|  | Succinic acid (%) | 0.04 ± 0 | 0.06 ± 0.01 | 0.04 ± 0 | 0.11 ± 0.03 | 0.08 ± 0.02 |
|  | **Total** | **3.22 ± 0.05** | **3.98 ± 0.04** | **3.13 ± 0.08** | **4.13 ± 0.16** | **4.83 ± 0.18** |
| **Sterol** |  |  |  |  |  |  |
|  | Campesterol (%) | 0.86 ± 0.04 | 0.59 ± 0.03 | 0.66 ± 0.04 | 0.52 ± 0.03 | 0.58 ± 0.05 |
|  | Cholesterol (%) | 34.97 ± 0.39 | 29.17 ± 1 | 35.32 ± 1.34 | 20.74 ± 3.38 | 27.43 ± 2.76 |
|  | Stigmast-5-ene (%) | 0.78 ± 0.01 | 0.59 ± 0 | 0.66 ± 0.03 | 0.53 ± 0.06 | 0.56 ± 0.05 |
|  | Stigmasterol (%) | 0.12 ± 0.01 | 0 ± 0 | 0 ± 0 | 0 ± 0 | 0.07 ± 0.06 |
|  | **Total** | **36.74 ± 0.11** | **30.35 ± 0.26** | **36.63 ± 0.35** | **21.78 ± 0.87** | **28.63 ± 0.73** |
| **Others** |  |  |  |  |  |  |
|  | 9-Octadecenamide (%) | 0.47 ± 0.01 | 0.49 ± 0.05 | 0.37 ± 0.12 | 0.44 ± 0.15 | 0.54 ± 0.2 |
|  | Decanamide (%) | 0.17 ± 0.01 | 0.13 ± 0.02 | 0.19 ± 0.02 | 0.14 ± 0.01 | 0.14 ± 0.01 |
|  | Ethanamine (%) | 0.13 ± 0.06 | 0.15 ± 0.09 | 0.1 ± 0.02 | 0.17 ± 0.05 | 0.16 ± 0.04 |
|  | Inosine (%) | 0.77 ± 0.04 | 0.96 ± 0.08 | 0.7 ± 0.02 | 1.12 ± 0.27 | 1.09 ± 0.15 |
|  | Uracil (%) | 0.07 ± 0 | 0.08 ± 0.01 | 0.06 ± 0.01 | 0.1 ± 0.04 | 0.09 ± 0.03 |
|  | Urea (%) | 0.54 ± 0.11 | 0.57 ± 0.03 | 0.44 ± 0.07 | 0.73 ± 0.24 | 0.86 ± 0.31 |
|  | **Total** | **2.15 ± 0.04** | **2.37 ± 0.05** | **1.86 ± 0.04** | **2.71 ± 0.13** | **2.88 ± 0.13** |
|  | **Total** | **100** | **100** | **100** | **100** | **100** |

**Table 9:** Compounds identification from the hexane extracts of whiteleg shrimp (*P. vannamei*) fed dietary treatments after 55 days. Values are presented in relative abundance (%) (means ± SD (n = 6)). P-values from Kruskal-Wallis (p ≤ 0.05).

| Identified compounds |  | Dietary treatments |  |  |  |  |
| --- | --- | --- | --- | --- | --- | --- |
|  |  | 55 Days |  |  |  |  |
|  |  | CTRL | SS5 | SS10 | SL5 | SL10 |
| **Alcohol** |  |  |  |  |  |  |
|  | Glycerol (%) | 0.22 ± 0.01 | 0.19 ± 0 | 0.14 ± 0.02 | 0.26 ± 0.02 | 0.15 ± 0.01 |
|  | 1-Hexadecanol (%) | 0.04 ± 0 | 0.04 ± 0 | 0.08 ± 0.01 | 0.05 ± 0 | 0.06 ± 0.04 |
|  | 9-Octadecen-1-ol (%) | 0.02 ± 0.02 | 0.06 ± 0 | 0.06 ± 0.01 | 0.08 ± 0.01 | 0.03 ± 0 |
|  | 1-Octadecanol (%) | 0.03 ± 0 | 0.02 ± 0 | 0.12 ± 0 | 0.04 ± 0 | 0.01 ± 0 |
|  | **Total** | **0.3 ± 0.006** | **0.31 ± 0.002** | **0.4 ± 0.01** | **0.42 ± 0.01** | **0.25 ± 0.013** |
| **Amino acid** |  |  |  |  |  |  |
|  | Alanine (%) | 3.06 ± 0.39 | 1.08 ± 0.08 | 1.29 ± 0.03 | 0.47 ± 0.07 | 7.33 ± 0.74 |
|  | Glycine (%) | 20.06 ± 2.2 | 18.39 ± 0.44 | 32.15 ± 0.82 | 3.15 ± 0.63 | 27.96 ± 1.76 |
|  | Isoleucine (%) | 0.38 ± 0.13 | 0.26 ± 0.04 | 0.54 ± 0.07 | 0.25 ± 0.01 | 0.29 ± 0.15 |
|  | Leucine (%) | 0.71 ± 0.23 | 0.53 ± 0.08 | 1.09 ± 0.12 | 0.5 ± 0.04 | 0.59 ± 0.3 |
|  | Ornithine (%) | 0.06 ± 0.01 | 0.15 ± 0.01 | 2.33 ± 0.29 | 0.06 ± 0.02 | 1.33 ± 0.14 |
|  | Phenylalanine (%) | 0.57 ± 0.06 | 0.36 ± 0.02 | 1.83 ± 0.09 | 0.2 ± 0.03 | 1.94 ± 0.09 |
|  | Proline (%) | 2.81 ± 0.18 | 1.1 ± 0.09 | 2.36 ± 0.06 | 1.4 ± 0.15 | 3.44 ± 0.22 |
|  | Threonine (%) | 0.1 ± 0.02 | 0.05 ± 0.01 |  | 0.05 ± 0.01 | 0.16 ± 0.02 |
|  | Tyrosine (%) | 0.52 ± 0.04 | 0.13 ± 0.01 | 0.8 ± 0.04 | 0.04 ± 0.03 | 2.87 ± 0.19 |
|  | Valine (%) | 2.1 ± 0.19 | 1.09 ± 0.08 | 2.12 ± 0.07 | 0.69 ± 0.08 | 2.79 ± 0.21 |
|  | **Total** | **30.37 ± 0.38** | **23.14 ± 0.12** | **44.51 ± 0.21** | **6.81 ± 0.13** | **48.69 ± 0.5** |
| **Fatty acid** |  |  |  |  |  |  |
|  | 11.14-Eicosadienoic acid (%) | 1.02 ± 0.01 | 1.33 ± 0.02 | 1.01 ± 0.01 | 1.76 ± 0.04 | 0.84 ± 0.01 |
|  | 11-Eicosaenoic acid (%) | 1.38 ± 0.04 | 1.32 ± 0.02 | 1.31 ± 0.03 | 1.91 ± 0.05 | 1.15 ± 0.01 |
|  | 9.12-Octadecadienoic acid (%) | 7.24 ± 0.27 | 8.53 ± 0.07 | 7.11 ± 0.08 | 11.13 ± 0.35 | 5.73 ± 0.05 |
|  | 9-Hexadecenoic acid (%) | 0.75 ± 0 | 0.9 ± 0.01 | 0.68 ± 0.01 | 1.25 ± 0.01 | 0.69 ± 0.02 |
|  | Arachidonic acid (%) | 0.39 ± 0.01 | 0.73 ± 0.01 | 0.44 ± 0.01 | 0.85 ± 0.03 | 0.58 ± 0.01 |
|  | Doconexent (%) | 0.86 ± 0.02 | 2.36 ± 0.03 | 1.17 ± 0.03 | 2.62 ± 0.08 | 0.7 ± 0.01 |
|  | Eicosanoic acid (%) | 0.27 ± 0 | 0.36 ± 0.01 | 0.25 ± 0.01 | 0.38 ± 0.01 |  |
|  | Eicosapentaenoic acid (%) | 2.16 ± 0.06 | 3.93 ± 0.04 | 2.26 ± 0.04 | 4.51 ± 0.17 | 1.58 ± 0.02 |
|  | Heptadecanoic acid (%) | 1.09 ± 0.01 | 1.31 ± 0.03 | 0.89 ± 0.02 | 1.5 ± 0.02 | 0.93 ± 0.02 |
|  | Myristic acid (%) | 0.33 ± 0.01 | 0.47 ± 0.01 | 0.43 ± 0.04 | 0.5 ± 0.01 | 0.44 ± 0.01 |
|  | Nonadecanoic acid (%) | 0.17 ± 0.01 | 0.24 ± 0.01 | 0.18 ± 0.01 | 0.28 ± 0.02 | 0.17 ± 0.01 |
|  | Oleic acid (%) | 11.25 ± 0.24 | 11.18 ± 0.05 | 9.61 ± 0.07 | 14.98 ± 0.18 | 8.44 ± 0.05 |
|  | Palmitic Acid (%) | 13.85 ± 0.53 | 14.41 ± 0.58 | 9.85 ± 0.42 | 18.11 ± 0.27 | 10.23 ± 0.39 |
|  | Pentadecanoic acid (%) | 0.24 ± 0 | 0.4 ± 0.01 | 0.23 ± 0.01 | 0.39 ± 0.01 | 0.25 ± 0.01 |
|  | Stearic acid (%) | 8.2 ± 0.45 | 8.81 ± 0.41 | 5.82 ± 0.28 | 10.67 ± 0.13 | 6.09 ± 0.25 |
|  | **Total** | **49.2 ± 0.1** | **56.3 ± 0.07** | **41.23 ± 0.07** | **70.85 ± 0.1** | **37.82 ± 0.06** |
| **Organic acid** |  |  |  |  |  |  |
|  | Aminomalonic acid (%) | 0.78 ± 0.03 | 0.53 ± 0.02 | 0.44 ± 0.06 | 0.97 ± 0.09 | 0.42 ± 0.03 |
|  | Behenic acid (%) | 0.13 ± 0 | 0.2 ± 0.01 | 0.14 ± 0.01 | 0.19 ± 0 | 0.14 ± 0.01 |
|  | Lactic Acid (%) | 0.5 ± 0.01 | 0.35 ± 0.03 | 0.29 ± 0.03 | 0.8 ± 0.06 | 0.36 ± 0.02 |
|  | Phosphoric acid (%) | 0.06 ± 0.01 | 0.08 ± 0.01 |  | 0.07 ± 0.05 | 0.06 ± 0 |
|  | Succinic acid (%) | 0.14 ± 0 |  |  |  | 0.25 ± 0.02 |
|  | Pentanedioic acid (%) | 0.31 ± 0 | 0.25 ± 0.01 | 0.41 ± 0.04 | 0.29 ± 0.07 | 0.36 ± 0.03 |
|  | **Total** | **1.92 ± 0.01** | **1.42 ± 0.02** | **1.28 ± 0.04** | **2.32 ± 0.06** | **1.59 ± 0.02** |
| **Sterol** |  |  |  |  |  |  |
|  | Campesterol (%) | 0.48 ± 0 | 0.46 ± 0.01 | 0.29 ± 0.01 | 0.53 ± 0.02 | 0.28 ± 0.01 |
|  | Cholesterol (%) | 17.15 ± 0.5 | 17.83 ± 0.23 | 11.88 ± 0.47 | 18.58 ± 0.96 | 11.11 ± 0.21 |
|  | Stigmast-5-ene (%) |  | 0.42 ± 0.01 | 0.36 ± 0.02 | 0.47 ± 0.03 | 0.26 ± 0.01 |
|  | **Total** | **18.05 ± 0.17** | **18.72 ± 0.08** | **12.53 ± 0.16** | **19.59 ± 0.34** | **11.66 ± 0.08** |
| **Others** |  |  |  |  |  |  |
|  | 9-Octadecenamide (%) | 0.09 ± 0.01 | 0.11 ± 0 | 0.04 ± 0.05 |  |  |
|  | Hexadecanamide (%) | 0.08 ± 0 |  |  |  |  |
|  | **Total** | **0.16 ± 0** | **0.11 ± 0** | **0.04 ± 0.05** |  |  |
| **Total** |  | 100 | 100 | 100 | 100 | 100 |

Looking at the sample extraction and analysis process, and since glycine is a 1) the possibility that this compound was not correctly extracted during the hexane extraction process, thus leading to a decrease in its abundance in the extract; 2) the silylation reaction does not occur completely and does not allow the derivatization of glycine and therefore, when injected into the GC-MS, it does not volatilize. The relative abundance of fatty acids is significantly higher and lower in the SL10 and SL5 groups, respectively, in relation to the CTRL. In sterols, the total abundance is higher in the SL5 group and lower in the SL10 group, compared to CTRL. Organic acids are significantly lower in the groups fed with stems. It is also important to note that the compounds identified after 31 days are the same as those identified after 55 days. The difference in the number of compounds identified in these two situations is related to the sensitivity of the equipment used, since these compounds are present in relative abundances of less than 1. Finally, it is verified that the relative abundance of nutritionally important compounds, such as amino acids and fatty acids, are not negatively affected by the incorporation of *S. ramosissima* in the feed of aquaculture animals.

#### 3.1.3. Haemocytes count

No differences in total haemocytes counts were observed during the feeding trial (Fig.1A). In response to the bath bacterial challenge with *Vibrio parahaemolyticus*, shrimp fed all experimental diets decreased the number of haemocytes in circulation following exposure to the pathogen, nonetheless no differences were observed between the experimental diets (Fig.1B).

**Figure 1.**
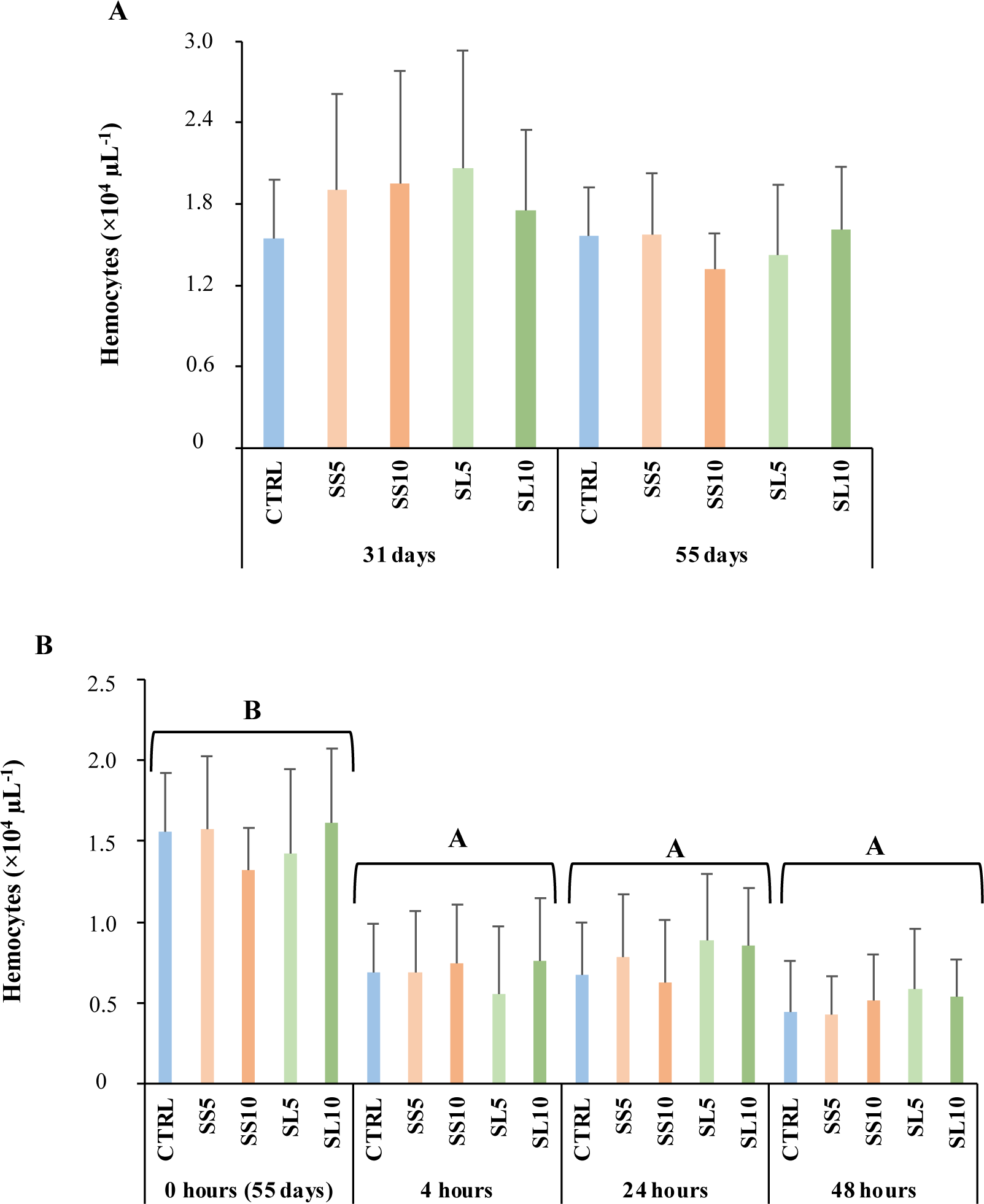
Haemocytes concentration during the feeding trial (A) and during the bacterial challenge (B) in whiteleg shrimp (*P. vannamei*). Experimental groups concern: control (Ctrl), fed with standard feed, and diets with *S. ramosissima* inclusion (SS5, SS10, SL5, and SL10). Values are means ± standard deviation (n=12). P-values from Two-way ANOVA (p ≤ 0.05). Tukey post-hoc test was used to identify differences in the experimental treatments in each sampling point. Different lowercase letters stand for significant differences between dietary treatments for the same time.

#### 3.1.4. Plasma humoral parameters

After 31 days of being fed experimental diets, shrimp that were given SS5, SL5, and SL10 exhibited lower levels of nitric oxide compared to those fed the CTRL diet. Similarly, shrimp fed SS5, SL5, and SL10 also showed lower lysozyme activity after 31 days, in contrast to those on the CTRL diet. The highest lysozyme activity was observed in shrimp that were fed a diet enriched with 10 % *S. ramosissima* stems (refer to Fig. 2 and Table S1). However, no significant differences were observed after 55 days of being fed the experimental diets (refer to Table S1 and Fig. 2). The complete set of results is present in the table S1 of the supplementary files.

**Figure 2.**
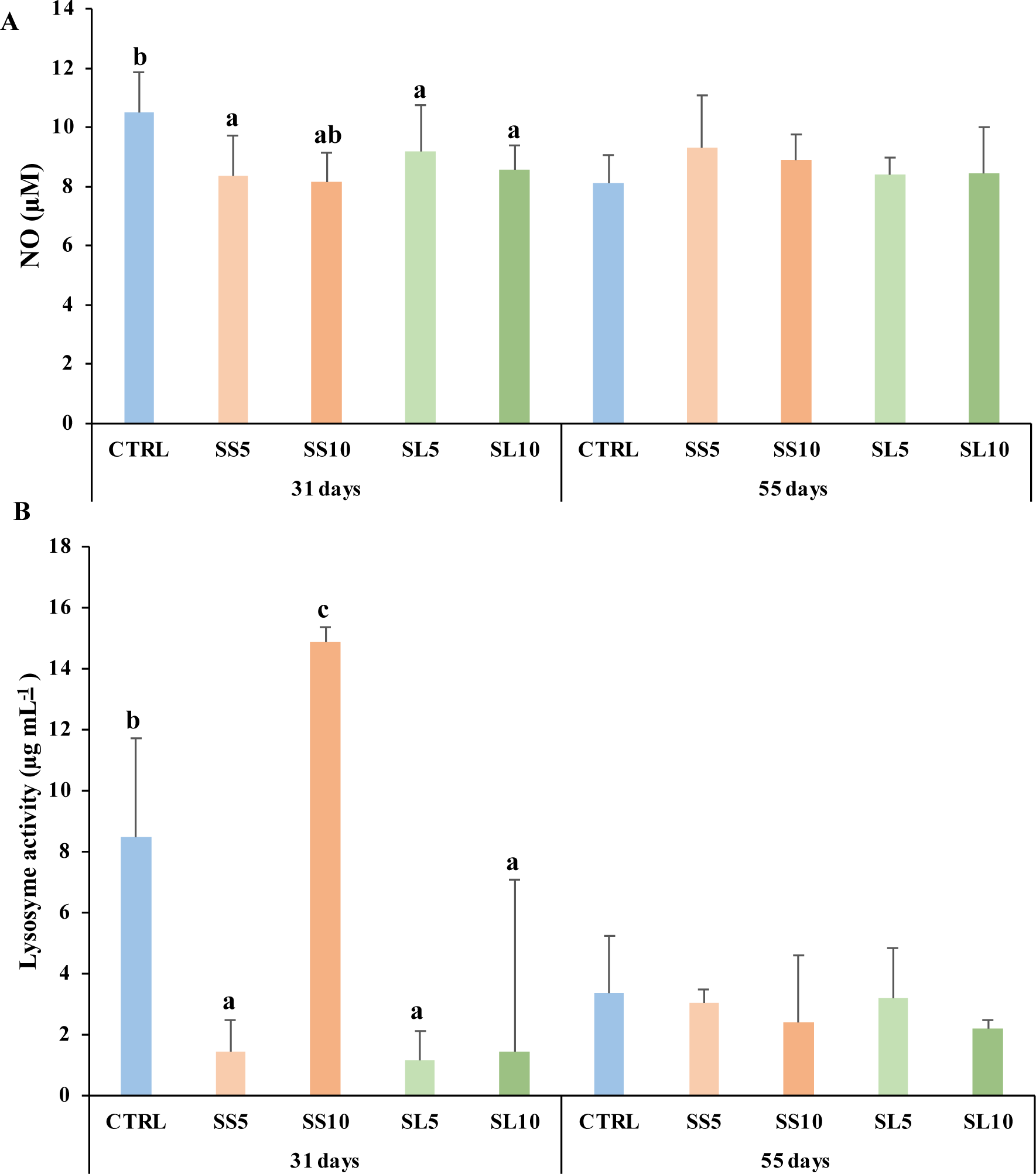
Plasma (A) nitric oxide (NO) and (B) lysozyme activity in whiteleg shrimp (*P. vannamei*) during the feeding trial. Experimental groups concern: control (Ctrl), fed with standard feed, and diets with *S. ramosissima* inclusion (SS5, SS10, SL5, and SL10). Values are means ± standard deviation (n=12). P-values from One-way ANOVA (p ≤ 0.05). Tukey post-hoc test was used to identify differences in the experimental treatments in each sampling point. Different lowercase letters stand for significant differences between dietary treatments for the same time.

#### 3.1.5. Hepatopancreas oxidant status

SOD and CAT activities displayed similar response profiles after 31 days of feeding trial. In whiteleg shrimp fed with stems, SOD and CAT activities decreased significantly in comparison with the CTRL group, with the exception of SL5 (Figs. 3A and 3B, respectively). Differently, at the end of the trial (day 55), a significant increase in SOD activity was observed in SS10 (Fig. 3A), and no significant variations in CAT activity were detected, in comparison with the CTRL group (Fig. 3B). Comparing the two levels leaves and seeds inclusion after 55 days, SL5 group showed lower CAT activity than SL10 (Fig. 3B).

**Fig. 3.**
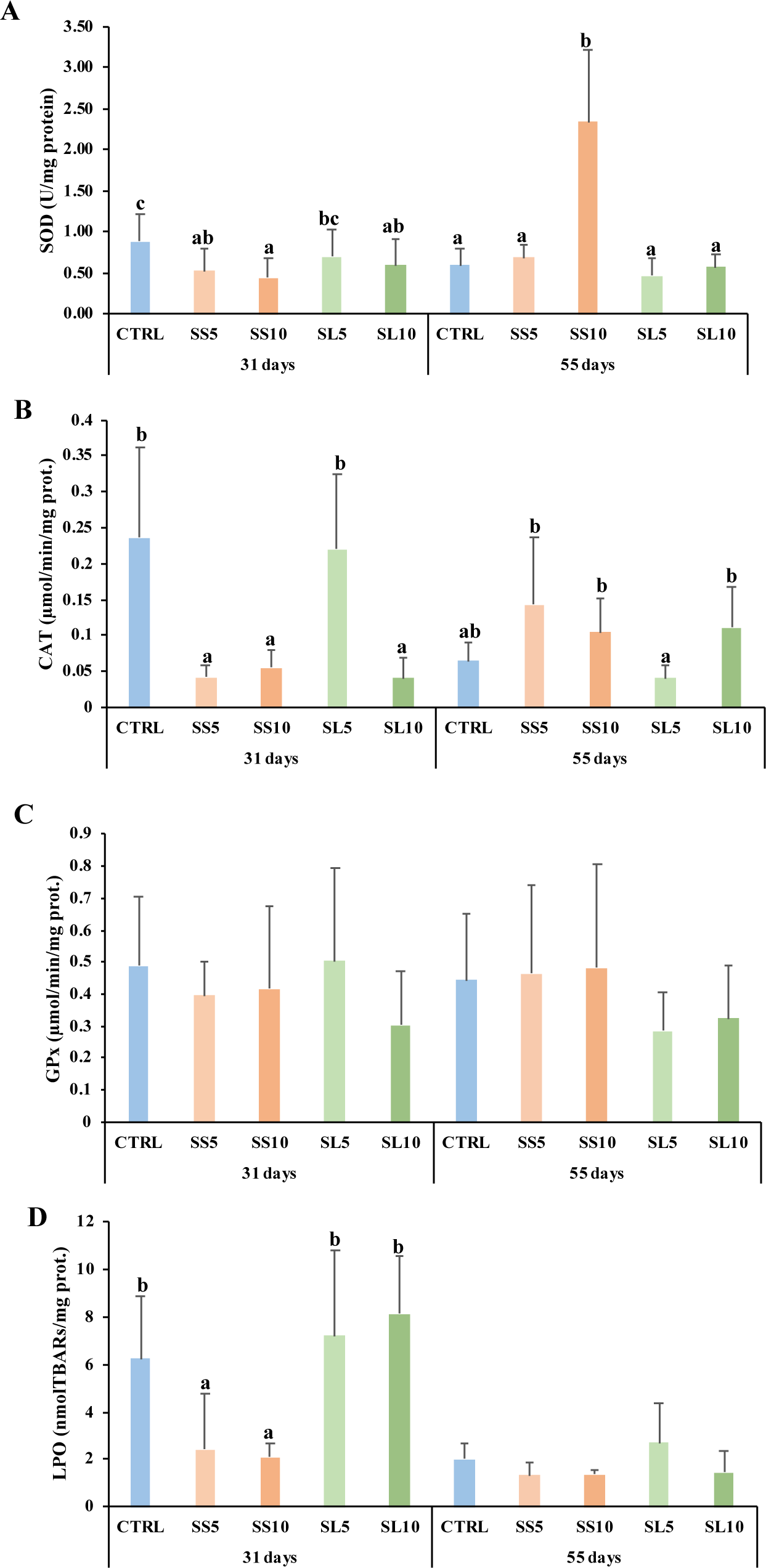
Mean values of hepatopancreas oxidative stress parameters in whiteleg shrimp (*P. vannamei*). (A) superoxide dismutase (SOD), (B) catalase (CAT), (C) glutathione peroxidase (GPx), (D) lipid peroxidation (LPO). Experimental groups concern: control (Ctrl), fed with standard feed, and diets with *S. ramosissima* inclusion (SS5, SS10, SL5, and SL10). Bars represent standard deviations. Different letters correspond to statistically significant differences (p<0.05).

GPx activity showed no significant variations, neither when compared with the CTRL group nor among diets with *S. ramosissima* inclusion (Fig. 3C).

Concerning LPO levels in hepatopancreas (Fig. 3D), significant alterations were restricted to the intermediate sampling (31 days), with both whiteleg shrimp groups fed with *S. ramosissima* stems (SS5 and SS10) displaying lower levels when compared to the CTRL group.

#### 3.1.6. Hepatopancreas gene expression

Regarding the health and antioxidant biomarkers analysed in shrimp hepatopancreas during the feeding period, main differences between dietary treatment were observed after 31 days of feeding (Table 10). On the one hand, it was observed a general downregulation of *casp3* in shrimp fed diets rich in *S. ramosissima* regardless the type or inclusion level. On the other hand, it was observed a general down-regulation of *gst* in shrimp fed diets rich in *S. ramosissima* regardless the type or inclusion level after 31 days of feeding. A downregulation of *gpx* was observed in shrimp fed SS10. After 55 days of feeding, a consistent increase in spermine synthase (SMS) was observed in shrimp that were fed diets rich in *S. ramosissima*, irrespective of the type or level of inclusion (Fig.4).

**Fig 4.**
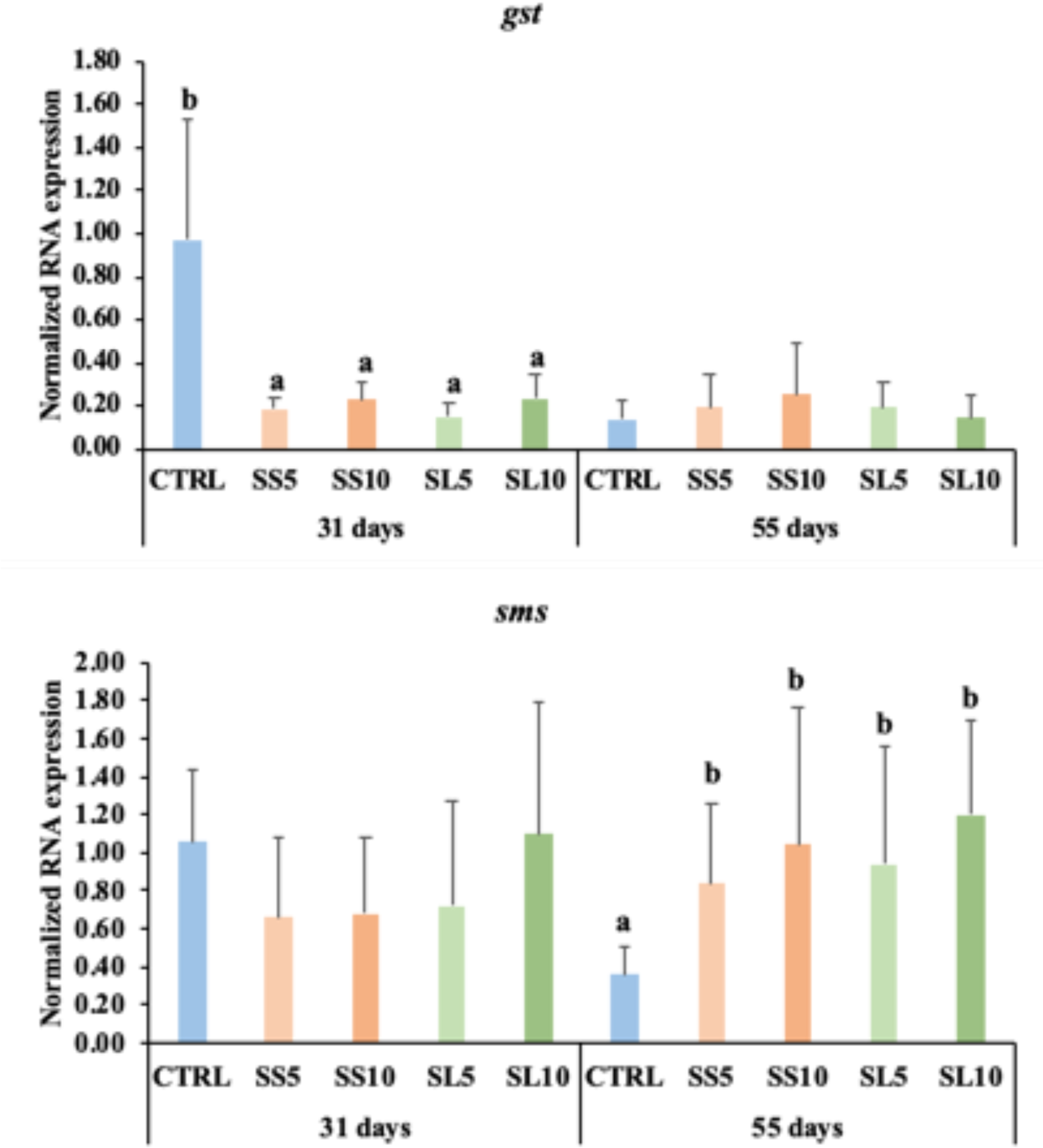
Relative expression of *gst* (Glutathione transferase) and *sms* (Spermine synthase) in the hepatopancreas gene expression of whiteleg shrimp (*P. vannamei*) fed dietary treatments during 55 days. Experimental groups concern: control (CTRL), fed with standard feed, and diets with *S. ramosissima* inclusion (SS5, SS10, SL5, and SL10). Values are presented as means ± SE (n = 12). P-values from One-way ANOVA (p ≤ 0.05). Tukey post-hoc test was used to identify differences in the experimental treatments in each sampling point. Different lowercase letters stand for significant differences between dietary treatments for the same time.

**Table 10.** Hepatopancreas gene expression of whiteleg shrimp (*P. vannamei*) fed dietary treatments during 31 and 55 days. Experimental groups concern: control (Ctrl), fed with standard feed, and diets with *S. ramosissima* inclusion (SS5, SS10, SL5, and SL10). Values are presented as means ± SD (n = 9). P-values from One-way ANOVA (p ≤ 0.05). Tukey post-hoc test was used to identify differences in the experimental treatments in each sampling point. Different lowercase letters stand for significant differences between dietary treatments for the same time. Different capital letters indicate differences between diets regardless of time or difference between times regardless of diets. Different symbols indicate difference between times for the same dietary treatment.

|  | Dietary treatments |  |  |  |  |  |  |  |  |  |
| --- | --- | --- | --- | --- | --- | --- | --- | --- | --- | --- |
|  | 31 Days |  |  |  |  | 55 Days |  |  |  |  |
|  | CTRL | SS5 | SS10 | SL5 | SL10 | CTRL | SS5 | SS10 | SL5 | SL10 |
| *pen3* | 1.61 ± 1.80 | 1.38 ± 0.48 | 1.57 ± 0.94 | 1.08 ± 0.39 | 2.47 ± 2.10 | 2.76 ± 1.28 | 2.87 ± 1.69 | 2.50 ± 1.96 | 4.30 ± 3.37 | 2.85 ± 1.15 |
| *lect* | 1.14 ± 0.52 | 1.18 ± 0.47 | 0.68 ± 0.28 | 1.30 ± 1.09 | 1.17 ± 0.43 | 1.39 ± 0.78 | 1.39 ± 0.32 | 0.82 ± 0.22 | 1.48 ± 0.81 | 1.01 ± 0.47 |
| *hmc* | 0.99 ± 0.51 | 0.67 ± 0.50 | 0.43 ± 0.19 | 1.24 ± 0.55 | 0.82 ± 0.32 | 0.32 ± 0.36 | 0.19 ± 0.09 | 0.35 ± 0.34 | 0.24 ± 0.10 | 0.32 ± 0.11 |
| *rab* | 1.19 ± 0.60 | 1.20 ± 0.61 | 1.14 ± 0.45 | 1.28 ± 0.86 | 1.61 ± 0.62 | 1.28 ± 0.91 | 1.58 ± 0.45 | 1.75 ± 0.45 | 1.56 ± 0.69 | 1.29 ± 0.57 |
| *lysC-like* | 1.02 ± 0.93 | 0.80 ± 0.73 | 0.44 ± 0.30 | 1.04 ± 0.78 | 0.91 ± 0.90 | 0.98 ± 0.71 | 0.76 ± 0.38 | 0.76 ± 0.80 | 0.51 ± 0.44 | 1.18 ± 1.12 |
| *gst* | 0.97 ± 0.57 | 0.18 ± 0.05 | 0.23 ± 0.09 | 0.15 ± 0.07 | 0.24 ± 0.11 | 0.14 ± 0.08 | 0.19 ± 0.16 | 0.25 ± 0.24 | 0.19 ± 0.12 | 0.14 ± 0.10 |
| *gpx* | 0.80 ± 0.27 | 0.58 ± 0.19 | 0.40 ± 0.13 | 0.65 ± 0.35 | 0.98 ± 0.46 | 0.76 ± 0.52 | 0.76 ± 0.26 | 0.73 ± 0.37 | 0.87 ± 0.43 | 0.85 ± 0.35 |
| *casp3* | 0.19 ± 0.08 | 0.05 ± 0.06 | 0.04 ± 0.03 | 0.09 ± 0.07 | 0.08 ± 0.06 | 0.01 ± 0.01 | 0.01 ± 0.00 | 0.02 ± 0.03 | 0.01 ± 0.01 | 0.01 ± 0.01 |
| *crus* | 1.33 ± 1.05 | 11.22 ± 20.74 | 1.72 ± 1.24 | 1.94 ± 1.16 | 13.19 ± 13.24 | 1.33 ± 0.80 | 1.93 ± 1.27 | 2.23 ± 1.10 | 2.99 ± 3.69 | 3.32 ± 2.66 |
| *trd* | 1.03 ± 0.60 | 1.09 ± 0.21 | 0.96 ± 0.43 | 0.85 ± 0.52 | 1.54 ± 0.55 | 1.09 ± 0.60 | 1.43 ± 0.49 | 1.15 ± 0.47 | 1.16 ± 0.32 | 1.51 ± 0.46 |
| *sms* | 1.06 ± 0.38 | 0.66 ± 0.42 | 0.68 ± 0.39 | 0.72 ± 0.56 | 1.10 ± 0.69 | 0.36 ± 0.14 | 0.83 ± 0.42 | 1.04 ± 0.72 | 0.94 ± 0.62 | 1.20 ± 0.49 |
| *sahh* | 1.19 ± 0.56 | 0.56 ± 0.35 | 1.01 ± 0.68 | 0.96 ± 0.49 | 1.06 ± 0.49 | 0.41 ± 0.28 | 1.04 ± 0.93 | 0.99 ± 0.77 | 0.88 ± 0.76 | 0.75 ± 0.60 |

|  |  | Dietary treatments |  |  |  |  |  |  |  |  |  |  |
| --- | --- | --- | --- | --- | --- | --- | --- | --- | --- | --- | --- | --- |
|  | One-way ANOVA | 31 Days |  |  |  |  | *p-value* | 55 Days |  |  |  |  |
|  | *p-value* | CTRL | SS5 | SS10 | SL5 | SL10 |  | CTRL | SS5 | SS10 | SL5 | SL10 |
| *pen3* | 0.482 | - | - | - | - | - | 0.796 | - | - | - | - | - |
| *lect* | 0.181 | - | - | - | - | - | 0.092 | - | - | - | - | - |
| *hmc* | 0.032 | **AB** | **AB** | **A** | **B** | **AB** | 0.652 | - | - | - | - | - |
| *rab* | 0.659 | - | - | - | - | - | 0.282 | - | - | - | - | - |
| *lysC-like* | 0.818 | - | - | - | - | - | 0.721 | - | - | - | - | - |
| *gst* | <0.001 | **B** | **A** | **A** | **A** | **A** | 0.831 | - | - | - | - | - |
| *gpx* | 0.012 | **B** | **AB** | **A** | **AB** | **B** | 0.827 | - | - | - | - | - |
| *casp3* | 0.005 | **B** | **A** | **A** | **A** | **A** | 0.085 | - | - | - | - | - |
| *crus* | 0.066 | - | - | - | - | - | 0.085 | - | - | - | - | - |
| *trd* | 0.159 | - | - | - | - | - | 0.348 | - | - | - | - | - |
| *sms* | 0.223 | - | - | - | - | - | 0.013 | **A** | **B** | **B** | **B** | **B** |
| *sahh* | 0.455 | - | - | - | - | - | 0.310 | - | - | - | - | - |

### 3.2. Bacterial challenge

#### 3.2.1. Plasma humoral parameters

Whiteleg shrimp fed SS5 and SL5 diets showed a higher production of nitric oxide than those fed the SS10 diet at 24 h after infection. ROS levels were higher in shrimps fed the SL10 diet when compared to those fed the SL5 after 48 h, in general this sampling point ROS levels were higher when compared to the previous samplings. Moreover, it was also observed a higher lysozyme activity in shrimp fed SL10 than those fed the other experimental diets at 48 h post infection (Fig. 5 and table S2). The complete set of results is present in the table S2 of the supplementary files.

**Figure 5.**
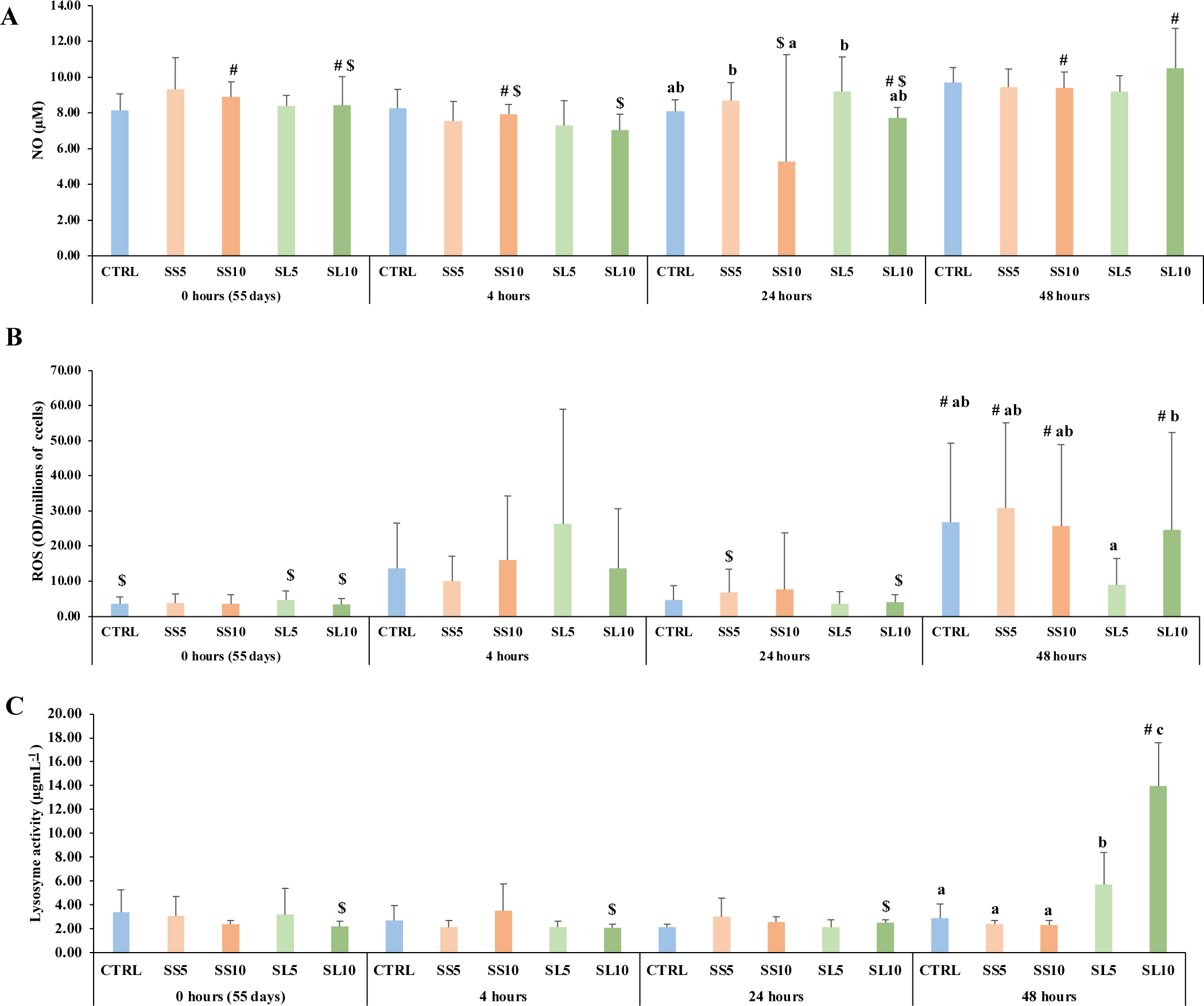
Plasma (A) nitric oxide (NO), (B) reactive oxygen species (ROS) levels and (C) lysozyme activity in whiteleg shrimp (*P. vannamei*) during the bacterial bath challenge. Experimental groups concern: control (Ctrl), fed with standard feed, and diets with *S. ramosissima* inclusion (SS5, SS10, SL5, and SL10). Values are means ± standard deviation (n=12). P-values from One-way ANOVA (p ≤ 0.05). Tukey post-hoc test was used to identify differences in the experimental treatments. Different lowercase letters stand for significant differences between dietary treatments for the same time. Different symbols indicate differences between times for the same dietary treatment.

#### 3.2.2. Hepatopancreas gene expression

The same panel of biomarkers studied during the feeding trial were also analysed for whiteleg shrimp sampled during the bacterial challenge. It was observed a general up-regulation of *sms* in shrimp fed diets rich in *S. ramosissima* regardless the type or inclusion level (Table 11). A down-regulation of *lys-like* was observed in shrimp fed SS10 and SL5 diets compared to those fed CTRL (Table 11). In contrast, an up-regulation of *pen3* was observed in shrimp fed SS10 regardless sampling time. Moreover, the expression of the *rab* was significantly up-regulated in the hepatopancreas of challenged whiteleg shrimp, firstly in those fed *S. ramosissima* stems (SS) at 5 % and 10 % after 4 h of infection and then of those fed *S. ramosissima* seeds and leaves (SL) at 5 % and 10 % inclusion levels after 48 h of bacterial challenge (Fig. 6).

**Figure 6.**
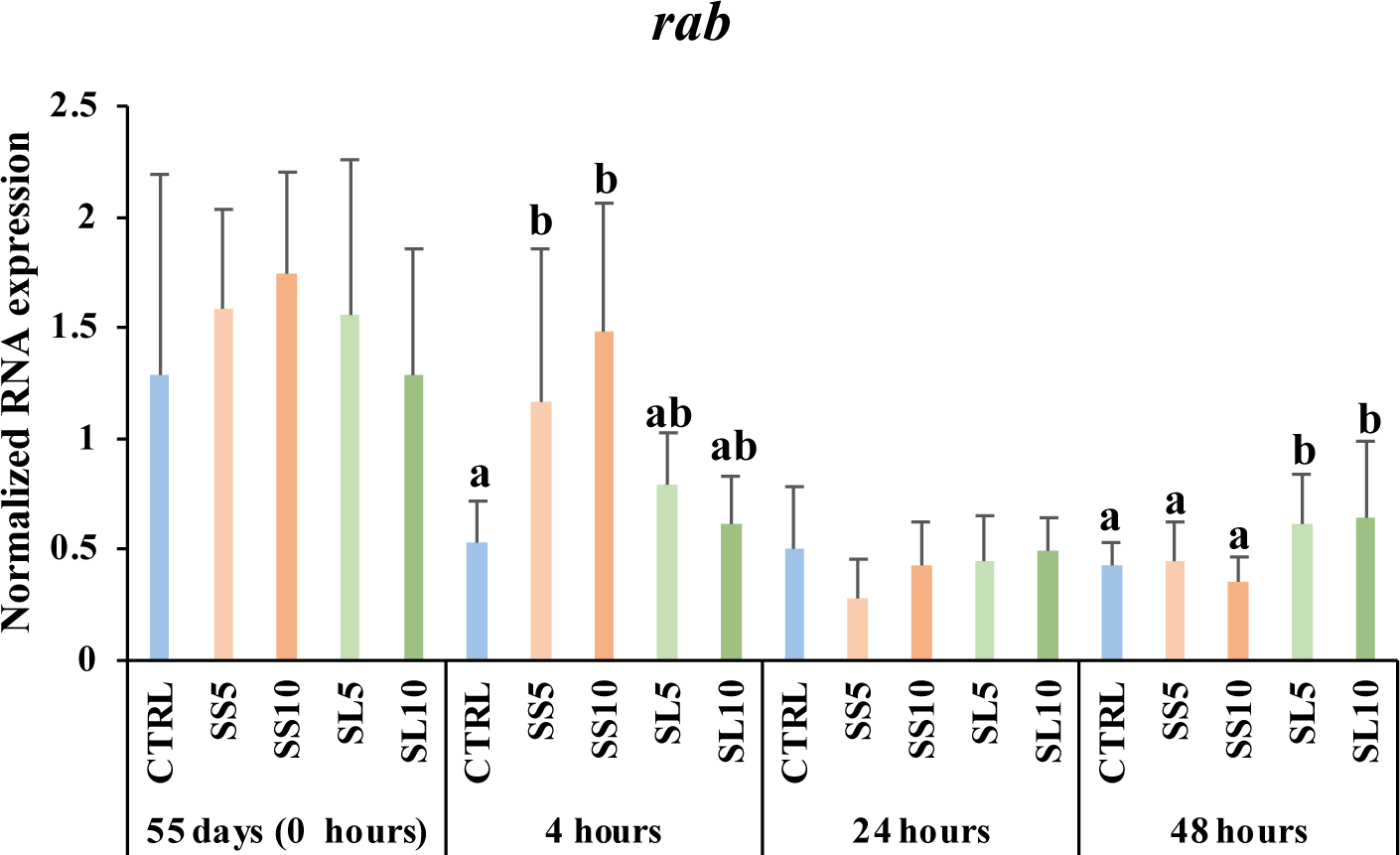
Relative expression of *rab* (Rab GTPase) in the hepatopancreas gene expression of whiteleg shrimp (*P. vannamei*) fed dietary treatments during the bath bacterial challenge. Experimental groups concern: control (Ctrl), fed with standard feed, and diets with *S. ramosissima* inclusion (SS5, SS10, SL5, and SL10). Values are presented as means ± SE (n = 12). P-values from Two-way ANOVA (p ≤ 0.05). Tukey post-hoc test was used to identify differences in the experimental treatments. Different lowercase letters stand for significant differences between dietary treatments for the same time.

**Table 11.**
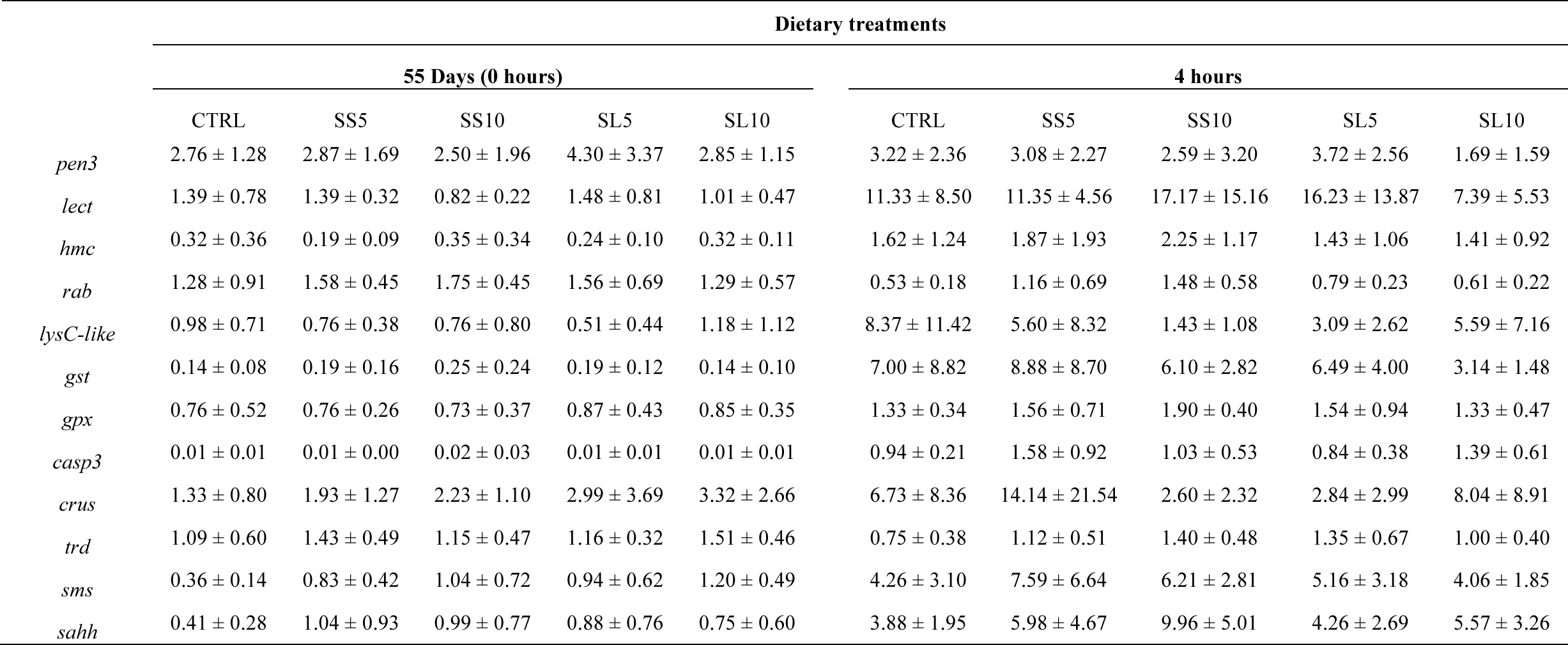
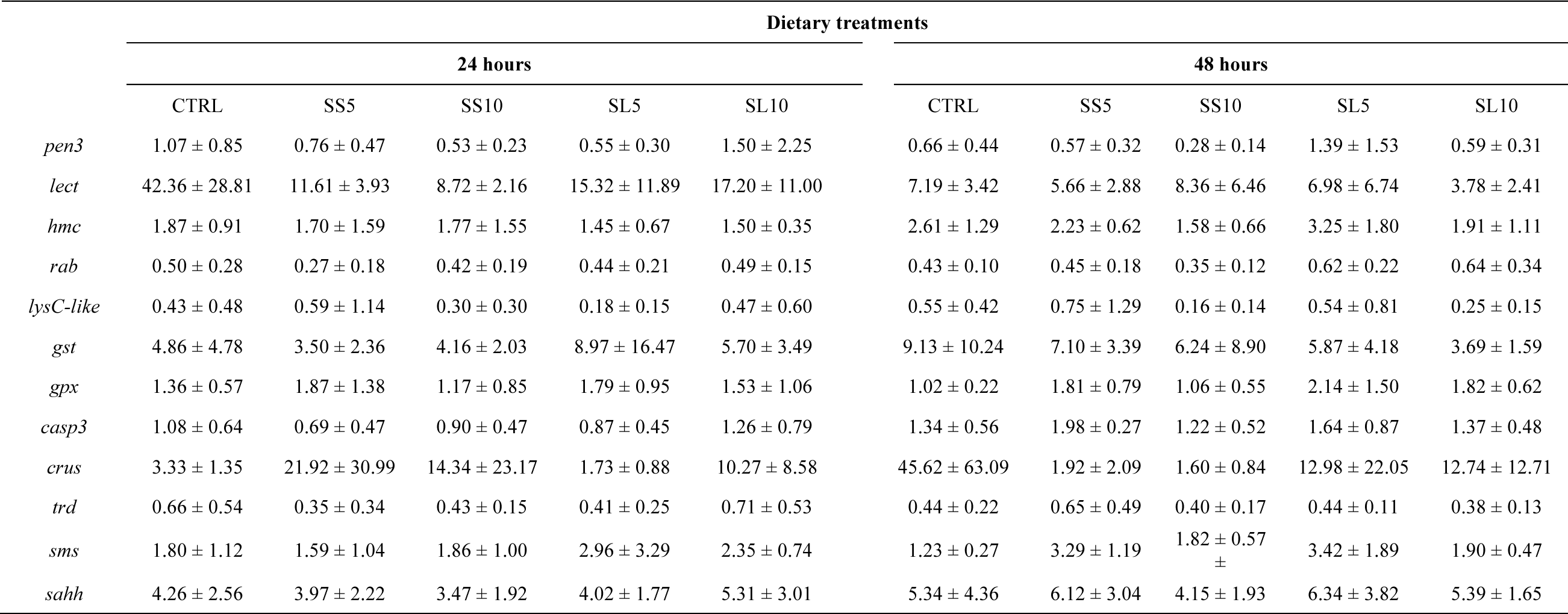
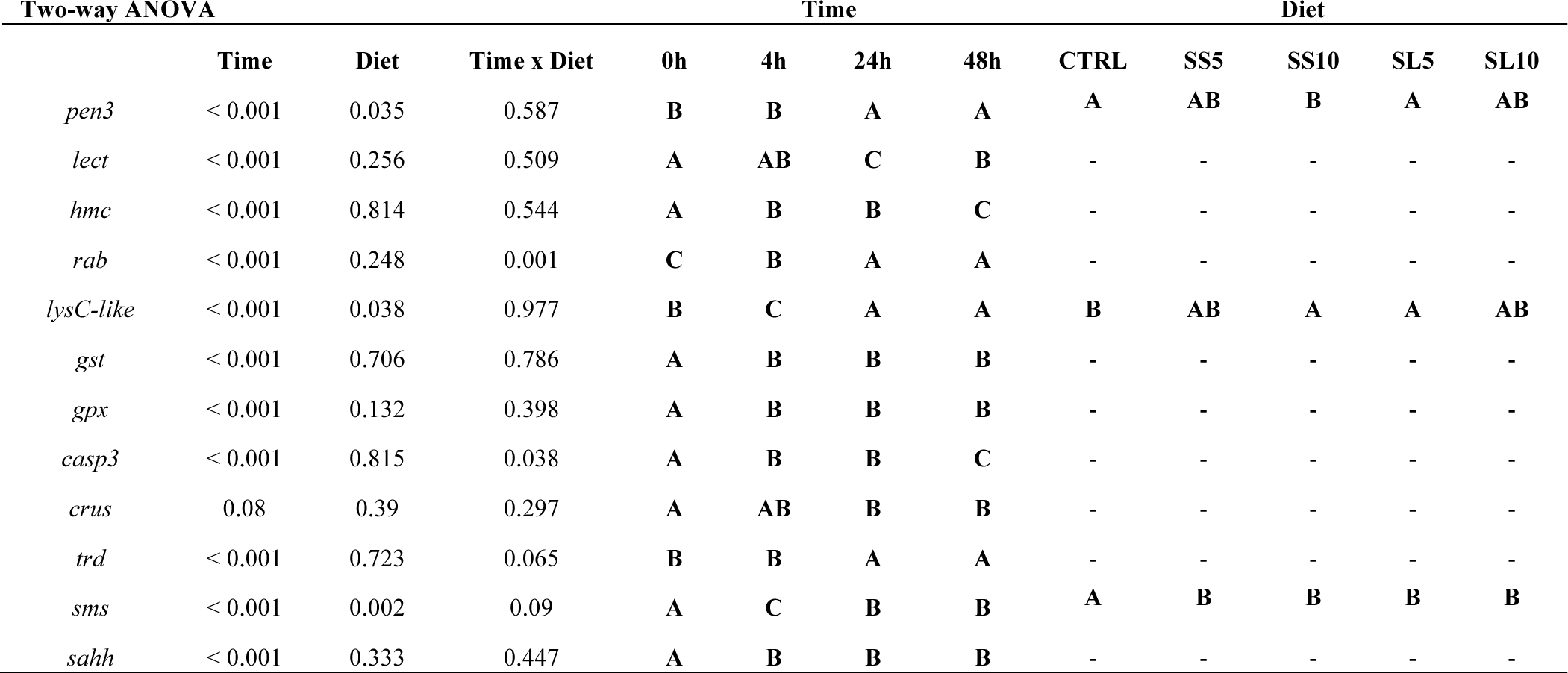
Hepatopancreas gene expression of whiteleg shrimp (*P. vannamei*) fed dietary treatments during the bath bacterial challenge. Experimental groups concern: control (Ctrl), fed with standard feed, and diets with *S. ramosissima* inclusion (SS5, SS10, SL5, and SL10). Values are presented as means ± SD (n = 9). P-values from Two-way ANOVA (p ≤ 0.05). Tukey post-hoc test was used to identify differences in the experimental treatments. Different lowercase letters stand for significant differences between dietary treatments for the same time. Different capital letters indicate differences between diets regardless of time or difference between times regardless of diets. Different symbols indicate differences between times for the same dietary treatment.

#### 3.2.3. Multivariate analysis from the hepatopancreas gene expression

An overall multivariate analysis combining raw data from hepatopancreas gene expression (using PCA-DA) was performed to discriminate the physiological effects caused by the experimental diets both at 31 and 55 days of feeding, and following the bacterial challenge (Fig. 7). The first two discriminant functions accounted for 81.79 % of dataset variability for feeding trial (31 and 55 days). Group discrimination was significant (Wilk’s lambda = 0.018, p <0.0001) highlighting the differences at the first sampling point (31 days), between CTRL and the experimental groups (p <0.001). While at the final sampling point, only differences between CTRL and SL10 and SS5 diets we observed. This discrimination was loaded by the expression of *hmc*, *casp3, gst*, and *gpx* (Fig. 7a).

**Figure 7.**
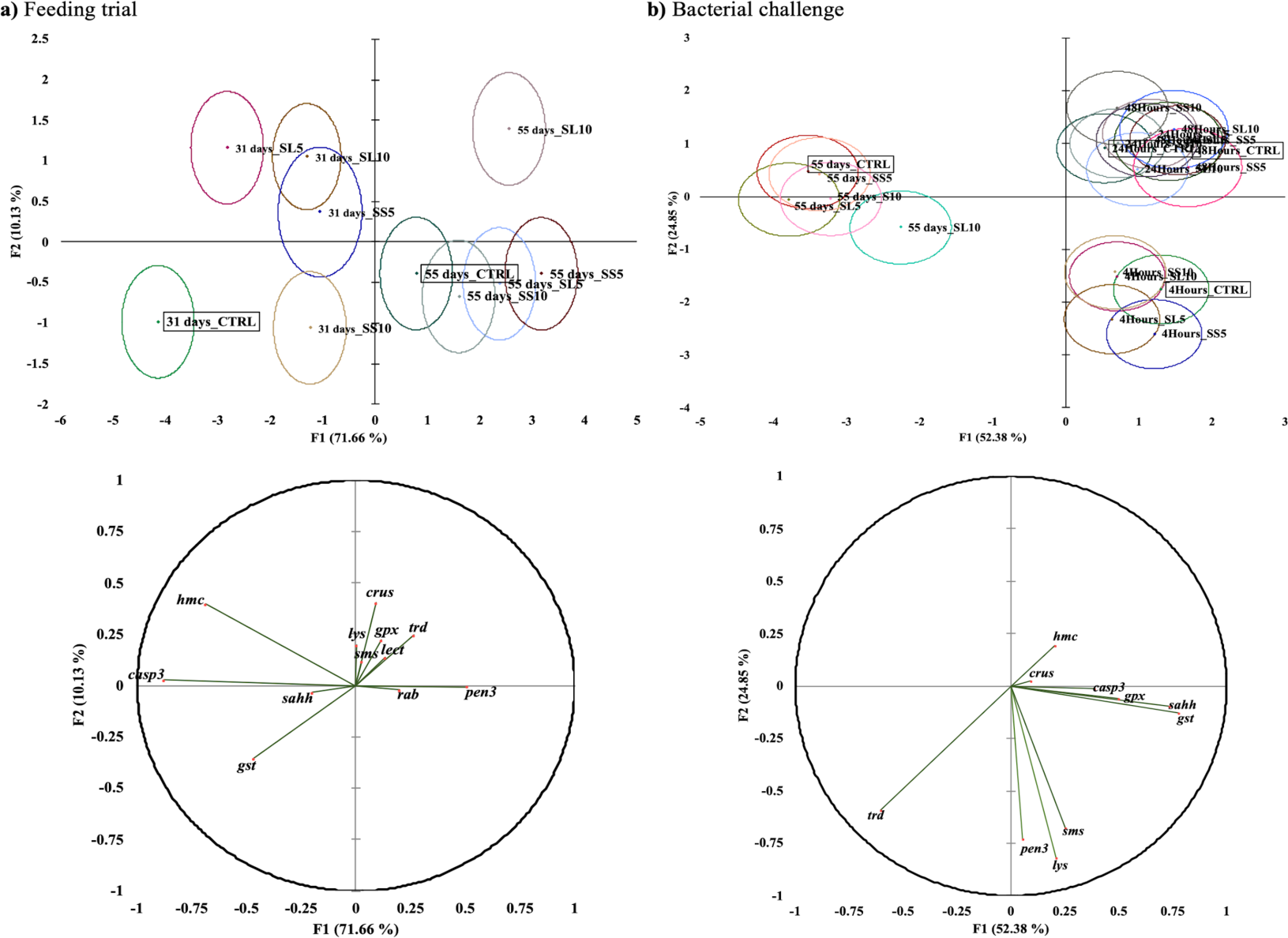
Canonical discriminant analysis of whiteleg shrimp (*P. vannamei*) immune and antioxidant responses biomarkers expression when fed with diets enriched with *S. ramosissima* during 31 and 55 days (a, feeding trial). And upon a bacterial challenge [b] (4, 24 and 48 h). Small dot marks represent group centroids, whereas ellipses indicate data distribution per group, and correlation variables/factors (factor loads) for two main discriminant functions (F1 and F2).

Following the bacterial challenge, groups were discriminated (Wilk’s lambda = 0.015; p <0.0001) and the first two discriminant factors accounted for 77.22 % of dataset variability. Overall, it is possible to observe a clear significant discriminant difference between sampling points regardless of dietary treatment. Particularly between the end of the feeding trial (prior to bacterial challenge) and 4 h following bacterial bath challenge (p <0.0001), and this separation was loaded by the expression of *trd*, *pen3*, *sms,* and *lys* (Fig. 7b).

## 4. Discussion

Recently, there has been a growing interest in utilizing halophytes as a healthy dietary option*. S. ramosissima* is one such halophyte, rich in bioactive compounds. A prominent feature of *S. ramosissima* is its high content of naturally occurring polyphenolic compounds, which are potent antioxidants with several health benefits. *S. ramosissima’s* polyphenolic profile includes flavonoids, tannins, and phenolic acids, all exhibiting various biological activities, including anti-inflammatory and antioxidant properties. Moreover, *S. ramosissima* is also a source of carotenoids, vitamins, and minerals, further contributing to its potential health-promoting effects ^35^. Hoping to understand the influence of incorporating *S. ramosissima* non-food fraction biomass in diets for the whiteleg shrimp regarding its effects on growth performance, survival, immune function, and oxidative status, shrimp juveniles were fed the experimental diets with *S. ramosissima* inclusion for 31 and 55 days. Subsequently, whiteleg shrimp were subjected to a bacterial bath challenge to assess their immune response when exposed to a pathogen. To the best of our knowledge, it is the first time that the inclusion of *S. ramosissima* is tested in aquafeeds and proposed as a growth modulator or nutraceutical ingredient for whiteleg shrimp.

Upon scrutinizing the photochemical profile analyses, noteworthy disparities emerged. At the 31-day mark, groups fed with *S. ramosissima* stems, leaves, and seeds demonstrated a reduced relative abundance of alcohols and sterols compared to the control group. This variance, particularly in groups SS10, SS5, SL5, and SL10, is statistically significant for alcohols, and for sterols, it is notably distinct in groups SL5 and SL10. Organic acids and fatty acids manifested significant differences between the control and the SL10 group, with the latter showcasing higher organic acid content and lower fatty acid levels. Amino acids, in turn, displayed markedly higher relative abundances in groups SS5, SL5, and SL10 compared to the control. This could be beneficial for the overall health and growth of the shrimp, as amino acids are essential for protein synthesis. The 55-day assessment revealed a significant elevation in the total relative abundance of alcohols in the SS10 and SL5 groups as opposed to the control. Intriguingly, the SL5 group exhibited a striking reduction in amino acid relative abundance, chiefly attributed to a notable decrease in glycine levels. Glycine is an essential amino acid and a key component of many biological processes. The reduction prompts questions about the overall nutritional quality of this specific diet ^36^.

In fact, it was also observed that the inclusion of *S. ramosissima* biomass in the experimental diets had no effects on the whiteleg shrimp weight, RGR and survival values after 31 and 55 days of feeding. However, feeding efficiency was compromised as shrimp fed diets with *S. ramosissima* inclusion had significantly higher FCR and feed intake values than the control diet at the end of the feeding trial. In fact, this was perceived after only 31 days of feeding for diets with 10 % inclusion levels, both in stems and seeds and leaves form. Although all diets had similar proximate compositions, these results suggest that the inclusion of *S. ramosissima* in the diets may have resulted in differences in digestibility, bioavailability of nutrients and metabolic utilisation by the shrimp. The inclusion of 10 % of *S. ramosissima* as seeds and leaves resulted in higher moisture levels in the shrimp whole body composition than at 5 % and when compared to the control diet. Despite not being supported by the statistical analysis, the SL10 diet tended to have lower dry matter contents, which can help explain these results. Furthermore, the ash levels in the shrimp whole body composition fed the SS10 diet was significantly higher than in those fed SL5, also reflecting the tendency to higher ash contents in the diets.

In the present study, feeds with *S. ramosissima* inclusion showed the ability to modulate the shrimp hepatopancreas antioxidant enzymes, specifically SOD and CAT enzyme activities. These enzymes cooperate in transforming toxic species in harmless water products, since SOD converts superoxide radicals into hydrogen peroxide and molecular oxygen, while CAT converts hydrogen peroxide into water ^37^. The most pronounced effects were visible after 31 days of feeding with SS (5 and 10 % inclusion levels) and with SL (10 % of inclusion). Moreover, *S. ramosissima* stems showed the potential to diminish the oxidative damage (measured as LPO), regardless of the dose incorporated. Interestingly, a down-regulation of the glutathione S-transferase gene (*gst*) was observed in shrimp fed diets rich in *S. ramosissima* regardless the biomass source or inclusion level after 31 days of feeding. Zhou, et al. ^38^ in a study with whiteleg shrimp, pointed out that GSTs are important components of various detoxification, antioxidant defence and stress-tolerance pathways. Therefore, the inclusion of *S. ramosissima* in aquafeeds for whiteleg shrimp juveniles may be a natural way of enhancing the animal’s health status. In fact, the gene expression of antioxidant enzymes (*gst* and *gpx*) indicates an overall profile of down-regulation. There seems to be an agreement between the general antioxidant profile at 31 days of feeding, measured both in activity and gene expression, which also aligns with the lipids peroxidation data (with emphasis on the reduction in SS). Overall, the data suggest that *S. ramosissima* dietary inclusion acts as an enhancer of the oxidant status, particularly when stems are incorporated and after 31 days of feeding. However, this effect appears to diminish after 55 days.

These results appear to confirm the contribution of non-enzymatic antioxidants by *S. ramosissima*, thereby sparing endogenous resources and creating a more favourable oxidant status. Despite the lack of studies in aquaculture species, the potential of *Salicornia* spp. as a functional feed capable of boosting the antioxidant response in farmed sheep has been demonstrated ^39,40^. In this study, higher FCR values were obtained with diets where *S. ramosissima* biomasses were included, which would increase the final production costs. However, potential antioxidant beneficial effects were observed when using the stems, which is considered a by-product of *S. ramosissima* cultivation and disposable. Therefore, by using a product that would otherwise be disposed instead of wheat meal, the price of the aquafeeds would be lower.

Moreover, the inclusion of *S. ramosissima* in shrimp diets was able to slightly modulate some of the plasma immune parameters and gene expression analysed on the hepatopancreas. No differences were observed in cell counts during the feeding trial, and a general decrease regardless of dietary treatments was observed following the bacterial bath challenge. Regarding the immune parameters analysed in haemolymph plasma, differences between dietary treatments were only observed after 31 days during the feeding trial, suggesting that the diet effect seems to disappear with a longer feeding period regime.

Decreases in plasma nitric oxide levels and lysozyme activity were observed in shrimp fed SS5, SL5 and SL10 after 31 days of feeding. However, whiteleg shrimp fed *S. ramosissima* stems (SS10) showed a higher lysozyme activity at that same time, suggesting a slightly increase of the immune response boosted by stems. Lysozyme is an enzyme with bacteriolytic properties, which can be found in both prokaryotes and eukaryotes. Its main function is to break down the β-1, 4-glycosidic bonds between *N-*acetylmuramic acid and *N-*acetylglucosamine in peptidoglycan, leading to bacterial lysis ^41^. It has been shown that lysozyme exhibits antimicrobial activity against both Gram-negative and Gram-positive bacteria, including Vibrio species that are harmful to shrimp ^42,43^. Interestingly, it was observed that shrimp fed *S. ramosissima* stems at 5 % and 10 % inclusion levels (SS5 and SS10, respectively) had higher levels of lysozyme activity after the bacterial challenge as well as respiratory burst (RB) particularly at 10 % inclusion level, and both after 48 h following bacterial bath challenge, suggesting that shrimp fed diets with *S. ramosissima* stems might be more prone to deal with the pathogen.

A mild modulation of gene expression during the feeding trial was observed, and particularly interesting to highlight is the up-regulation of *spermine synthase* after 55 days of feeding in all experimental diets with *S. ramosissima* regardless of the biomass type or the inclusion level. Spermine synthase is an enzyme that catalyses the conversion of spermidine to spermine, which are polyamines involved in various cellular processes such as cell growth, proliferation, and differentiation ^44^. Interestingly, Machado, et al. ^34^ observed a modulation of the methionine-related pathways with the up-regulation of hepatopancreas mRNA expression of SAM-synthase (SAM-synth) and also the spermine synthase (SMS) in shrimp fed a diet rich in AQUAVI® Met-Met pointing to stimulation of polyamine biosynthesis route and key for cell proliferation.

The expression of the rab (Rab GTPase) gene responsible for the formation, transport, and fusion of intracellular organelles crucial for immune response was higher in the hepatopancreas of challenged shrimp fed *S. ramosissima* stems (SS) at 5% and 10% inclusion levels already at 4 h after infection. The Rab protein is an important regulatory component involved in cell biosynthesis, endocytosis, and vesicle transport ^45^. Recent research has also demonstrated its essential role in the development and maturation of phagocytes and innate pathways that aid in the removal of foreign pathogens ^46^. These findings support the potential health-promoting benefits of Salicornia as mentioned earlier.

To gain a comprehensive understanding of the dietary impact on hepatopancreas gene expression, a multivariate analysis combining data from various genes was conducted. The results revealed that shrimp fed SL10 were markedly distinct from those fed the remaining experimental diets after 55 days of feeding. However, it was impossible to draw any firm conclusions from the bacterial challenge gene expression results, as no significant differences were observed among shrimp fed on different diets. Nevertheless, a clear effect of the bacterial bath challenge was evident, as demonstrated by a sharp distinction between shrimp sampled at 0 hours (55 days of feeding) and those collected after 4 hours of the bacterial bath. Furthermore, a distinct separation was noted between 4 hours and the cluster of 24- and 48-hours post-infection.

In summary, this study demonstrates that integrating *S. ramosissima* biomass into the diets of juvenile whiteleg shrimp does not compromise their growth, survival, or overall health. This breakthrough offers a substantial opportunity to reinforce halophyte production and advance sustainable aquafeed practices, potentially supplanting commonplace ingredients like wheat meal, a valuable resource also utilized in human consumption. Moreover, the study’s promising outcome underscores the potential antioxidant advantages stemming from the utilization of *S. ramosissima* stems, conventionally viewed as cultivation by-products often relegated to waste. This innovative strategy not only taps into an overlooked resource but also presents a cost-effective alternative to traditional aquafeed components like wheat meal. Despite a perceived impact on shrimp feeding efficiency, the incorporation of *S. ramosissima* biomass stands as a promising measure to mitigate the rising costs associated with cereal-based feeds. The cultivation of *S. ramosissima* is in line with the current global emphasis on resource optimization, waste reduction, and the exploration of alternative energy sources, positioning it as a viable element of a sustainable circular economy.

## 5. Ethics statement

All activities described were undertaken within the clear boundaries of national and EU legal frameworks, were directed by qualified scientists/technicians and were conducted according to the European guidelines on the protection of animals used for scientific purposes (Directive 2010/63/UE of the European Parliament and on the European Union Council) and under strict monitoring and control of DGAV—(Direção Geral de Alimentação e Veterinária), Animal Welfare Division, which is the competent authority responsible for implementing the legislation on the “protection of animals used for scientific purposes” in Portugal.

## 6. Author contribution

RJMR, MP and BC conceived the experiments and MM, LRP, AB and AL conducted the experimental trials. AL was responsible for the harvest of *S. ramosissima*. JD formulated and manufactured the experimental diets. LRP, MM, RM, SFB, CT, AC, SG, ACSV and DCGAP assisted with the analytical procedures. LRP wrote the manuscript under the supervision of MP and BC. All authors contributed to and approved the manuscript.

## 8. Data availability

The data that support the findings of this study are available from the corresponding author upon request.

## 9. Acknowledgements

The authors would like to acknowledge technical support from Bruna Silva during shrimp rearing and sampling at Riasearch; the SPAROS feed production team for manufacturing the tested microdiets, Diogo Peixoto and Paulo Santos from CIIMAR for their help during the bacterial challenge procedures. Also, the authors acknowledge support from Renata Serradeiro in funding acquisition and in the conceptualization of this study.

