## Supplementary material for "Inclusion of *Salicornia ramosissima* biomass in diets for juvenile whiteleg shrimp (*Penaeus vannamei*) induces favourable but transient effects in the immune and oxidative status": Supplemary files 1

[1]*Centro Interdisciplinar de Investigação Marinha e Ambiental (CIIMAR), Universidade do Porto, Terminal de Cruzeiros do Porto de Leixões, Avenida General Norton de Matos, S/N, 4450-208 Matosinhos, Portugal.*
[2]*Centre for Environmental and Marine Studies (CESAM) and Department of Biology, University of Aveiro, Santiago University Campus, 3810-193 Aveiro, Portugal.*
[3]*Riasearch Lda, Cais da Ribeira de Pardelhas nº21, Murtosa, Portugal.*
[4] *SPAROS Lda, Área Empresarial de Marim, Lote C, 8700-221 Olhão, Portugal.*
[5]*Instituto de Ciências Biomédicas Abel Salazar (ICBAS-UP), Universidade do Porto, Rua de Jorge Viterbo Ferreira nº 228, 4050-313 Porto, Portugal.*
[6]*LAQV-REQUIMTE, Department of Chemistry, Campus Universitário de Santiago, University of Aveiro, 3810-193 Aveiro, Portugal*

**Table S1**. Plasma humoral parameters of whiteleg shrimp (*P. vannamei*) fed dietary treatments during 31 and 55 days. Experimental groups concern: control (Ctrl), fed with standard feed, and Salicornia supplemented diets (SS5, SS10, SL5, and SL10). Values are presented as means ± SD (n = 12). P-values from One-way ANOVA ($p \leq 0.05$). Tukey post-hoc test was used to identify differences in the experimental treatments in each sampling point. Different lowercase letters stand for significant differences between dietary treatments for the same time.

| Parameters | Dietary treatments | | | | | |
|---|---|---|---|---|---|---|
| | 31 Days | | | | | One-Way ANOVA |
| | CTRL | SS5 | SS10 | SL5 | SL10 | *p-value* |
| Proteins (mg $mL^{-1}$) | 76.18 ± 16.31 | 79.95 ± 15.22 | 70.06 ± 24.17 | 72.24 ± 15.63 | 66.83 ± 10.92 | ns |
| Prophenoloxidase (U $mL^{-1}$) | 0.014 ± 0.008 | 0.019 ± 0.015 | 0.026 ± 0.015 | 0.010 ± 0.014 | 0.017 ± 0.012 | ns |
| Lysozyme (µg $mL^{-1}$) | 8.47 ± 3.24 $^{b}$ | 1.44 ± 0.94 $^{a}$ | 14.88 ± 5.66 $^{c}$ | 1.17 ± 0.49 $^{a}$ | 1.43 ± 1.05 $^{a}$ | <0.001 |
| NO (µM) | 10.50 ± 1.36 $^{b}$ | 8.36 ± 1.35 $^{a}$ | 8.16 ± 0.97$^{c}$ | 9.18 ± 1.56 $^{a}$ | 8.58 ± 0.83 $^{a}$ | 0.002 |
| Anti-Protease (%) | 12.36 ± 4.00 | 17.25 ± 7.35 | 16.12 ± 5.70 | 18.48 ± 4.63 | 14.55 ± 6.27 | ns |
| Protease (%) | 10.13 ± 0.32 | 10.21 ± 0.37 | 10.18 ± 0.25 | 14.00 ± 7.71 | 10.18 ± 0.26 | ns |
| Hemocyanin (mmol $L^{-1}$) | 0.22 ± 0.03 | 0.23 ± 0.04 | 0.22 ± 0.04 | 0.22 ± 0.03 | 0.21 ± 0.03 | ns |
| RB (OD/millions of cells) | 19.84 ± 16.73 | 17.36 ± 9.85 | 22.58 ± 14.07 | 27.05 ± 17.84 | 22.11 ± 9.48 | ns |

| Parameters | Dietary treatments | | | | | |
|---|---|---|---|---|---|---|
| | 55 Days | | | | | One-Way ANOVA |
| | CTRL | SS5 | SS10 | SL5 | SL10 | *p-value* |
| Proteins (mg $mL^{-1}$) | 77.67 ± 12.09 | 75.57 ± 12.30 | 81.49 ± 18.46 | 86.72 ± 16.32 | 85.21 ± 18.07 | ns |
| Prophenoloxidase (U $mL^{-1}$) | 0.009 ± 0.006 | 0.022 ± 0.012 | 0.024 ± 0.015 | 0.026 ± 0.021 | 0.017 ± 0.013 | ns |
| Lysozyme (µg $mL^{-1}$) | 3.36 ± 1.89 | 3.05 ± 1.65 | 2.39 ± 0.30 | 3.19 ± 2.22 | 2.20 ± 0.42 | ns |
| NO (µM) | 8.12 ± 0.95 | 9.31 ± 1.77 | 8.88 ± 0.86 | 8.41 ± 0.56 | 8.45 ± 1.57 | ns |
| Anti-Protease (%) | 21.00 ± 9.24 | 20.65 ± 4.90 | 21.38 ± 7.99 | 16.50 ± 6.63 | 22.90 ± 7.25 | ns |
| Protease (%) | 10.49 ± 0.37 | 10.42 ± 0.25 | 10.68 ± 0.60 | 12.34 ± 5.44 | 10.26 ± 0.23 | ns |
| Hemocyanin (mmol $L^{-1}$) | 0.22 ± 0.04 | 0.23 ± 0.03 | 0.24 ± 0.04 | 0.23 ± 0.03 | 0.21 ± 0.04 | ns |
| RB (OD/millions of cells) | 3.54 ± 2.02 | 3.79 ± 2.54 | 3.51 ± 2.72 | 4.72 ± 2.42 | 3.30 ± 1.83 | ns |

**Table S2.** Plasma innate immune parameters of whiteleg shrimp (*P. vannamei*) dietary treatments during the bacterial bath challenge. Experimental groups concern: control (Ctrl), fed with standard feed, and diets with *S. ramosissima* inclusion (SS5, SS10, SL5, and SL10). Values are presented as means ± SD (n = 12). P-values from One-way ANOVA ($p \leq 0.05$). Tukey post-hoc test was used to identify differences in the experimental treatments. Different lowercase letters stand for significant differences between dietary treatments for the same time. Different capital letters indicate differences between diets regardless of time or difference between times regardless of diets. Different symbols indicate differences between times for the same dietary treatment.

| **Parameters** | **Dietary treatments** | | | | | | | | | |
|---|---|---|---|---|---|---|---|---|---|---|
| | **55 Days (0 Hours)** | | | | | **4 hours** | | | | |
| | CTRL | SS5 | SS10 | SL5 | SL10 | CTRL | SS5 | SS10 | SL5 | SL10 |
| Proteins (mg/mL) | 77.67 ± 12.09 | 75.57 ± 12.30 | 81.49 ± 18.46 | 86.72 ± 16.32 | 85.21 ± 18.07 | 84.34 ± 22.42 $^{ab}$ | 71.55 ± 21.82 $^{ab}$ | 99.24 ± 14.82 $^{a}$ | 60.79 ± 20.99 $^{ab}$ | 58.91 ± 17.96 $^{b}$ |
| Prophenoloxidase (U/mL) | 0.009 ± 0.006 | 0.022 ± 0.012 | 0.024 ± 0.015 | 0.026 ± 0.021 | 0.017 ± 0.013 | 0.023 ± 0.020 | 0.022 ± 0.022 | 0.033 ± 0.024 | 0.028 ± 0.025 | 0.017 ± 0.024 |
| Lysozyme (µg/ml) | 3.36 ± 1.89 | 3.05 ± 1.65 | 2.39 ± 0.30 | 3.19 ± 2.22 | 2.20 ± 0.42$^{\$}$ | 2.72 ± 1.22 | 2.14 ± 0.57 | 3.51 ± 2.28 | 2.11 ± 0.53 | 2.07 ± 0.33$^{\$}$ |
| NO (µM) | 8.12 ± 0.95 | 9.31 ± 1.77 | 8.88 ± 0.86 $^{\#}$ | 8.41 ± 0.56 | 8.45 ± 1.57$^{\#\$}$ | 8.27 ± 1.03 | 7.53 ± 1.13 | 7.93 ± 0.54$^{\#\$}$ | 7.29 ± 1.40 | 7.05 ± 0.86$^{\$}$ |
| Hemocyanin (mmol/L) | 0.22 ± 0.04 | 0.23 ± 0.03 | 0.24 ± 0.04 | 0.23 ± 0.03 | 0.21 ± 0.04 | 0.20 ± 0.05 | 0.20 ± 0.03 | 0.22 ± 0.09 | 0.18 ± 0.05 | 0.19 ± 0.05 |
| RB (OD/millions of cells) | 3.54 ± 2.02$^{\$}$ | 3.79 ± 2.54 | 3.51 ± 2.72 | 4.72 ± 2.42$^{\$}$ | 3.30 ± 1.78$^{\$}$ | 13.67 ± 12.77 | 9.94 ± 7.15 | 16.06 ± 18.32 | 26.37 ± 32.66 | 13.57 ± 17.08 |

| **Parameters** | **Dietary treatments** | | | | | | | | | |
|---|---|---|---|---|---|---|---|---|---|---|
| | **24 hours** | | | | | **48 hours** | | | | |
| | CTRL | SS5 | SS10 | SL5 | SL10 | CTRL | SS5 | SS10 | SL5 | SL10 |
| Proteins (mg/mL) | 68.64 ± 22.62 | 75.25 ± 34.14 | 74.78 ± 18.71 | 68.78 ± 19.59 | 66.97 ± 16.28 | 65.83 ± 13.28 | 74.42 ± 27.96 | 69.45 ± 15.81 | 64.85 ± 21.27 | 54.77 ± 13.67 |
| Prophenoloxidase (U/mL) | 0.004 ± 0.007 | 0.030 ± 0.022 | 0.024 ± 0.017 | 0.014 ± 0.013 | 0.006 ± 0.006 | 0.017 ± 0.017 | 0.019 ± 0.018 | 0.024 ± 0.014 | 0.024 ± 0.025 | 0.026 ± 0.035 |
| Lysozyme (µg/ml) | 2.11 ± 0.29 | 3.00 ± 1.54 | 2.58 ± 0.40 | 2.13 ± 0.59 | 2.52 ± 0.22$^{\$}$ | 2.89 ± 1.20 | 2.38 ± 0.28 | 2.29 ± 0.38 | 5.71 ± 2.65 | 13.97 ± 3.65$^{\#}$ |
| NO (µM) | 8.09 ± 0.64$^{ab}$ | 8.68 ± 1.01$^{a}$ | 5.27 ± 5.98$^{\$b}$ | 9.19 ± 1.95$^{a}$ | 7.72 ± 0.57$^{\$\#ab}$ | 9.71 ± 0.82 | 9.45 ± 1.00 | 9.41 ± 0.88$^{\#}$ | 9.19 ± 0.87 | 10.49 ± 2.25$^{\#}$ |
| Hemocyanin (mmol/L) | 0.20 ± 0.04 | 0.23 ± 0.05 | 0.21 ± 0.04 | 0.21 ± 0.04 | 0.20 ± 0.04 | 0.18 ± 0.04 | 0.20 ± 0.04 | 0.20 ± 0.03 | 0.19 ± 0.04 | 0.18 ± 0.03 |
| RB (OD/millions of cells) | 4.65 ± 4.09 | 6.85 ± 6.68 | 7.75 ± 16.03 | 3.57 ± 3.51$^{\$}$ | 3.98 ± 2.12$^{\$}$ | 26.84 ± 22.49$^{\#ab}$ | 30.93 ± 24.23$^{\#\$b}$ | 25.75 ± 23.16$^{\#a}$ | 9.00 ± 7.45$^{\#ab}$ | 24.66 ± 27.65$^{\#ab}$ |

**Two-way ANOVA**

| Parameters | Time | Diet | Time x diet | Time | | | | Diet | | | | |
|---|---|---|---|---|---|---|---|---|---|---|---|---|
| | | | | 0h (55D) | 4 hours | 24 hours | 48 hours | CTRL | SS5 | SS10 | SL5 | SL10 |
| **Proteins (mg/mL)** | 0.0033 | 0.0384 | 0.0434 | B | AB | AB | AB | AB | AB | B | AB | A |
| **Pro-Fenoloxidase (U/mL)** | ns | ns | ns | - | - | - | - | - | - | - | - | - |
| **Lisozima (µg/ml)** | <0.001 | <0.001 | <0.001 | A | A | A | B | A | A | A | B | C |
| **NO (µM)** | <0.001 | ns | 0.0123 | AB | AB | AB | B | - | - | - | - | - |
| **Hemocianina (mmol/L)** | <0.001 | ns | ns | AB | AB | AB | A | - | - | - | - | - |
| **ROS (Abs/milhão de células)** | <0.001 | ns | 0.025 | A | AB | A | B | - | - | - | - | - |
